# Environmental temperature is a strong driver of subspecies competition in the *Drosophila* microbiome

**DOI:** 10.64898/2026.02.19.706767

**Authors:** Bosco Gracia-Alvira, Stefanie Migotti, Xiaomeng Tian, Viola Nolte, Christian Schlötterer

## Abstract

Most microbiome research focusses on the taxonomic composition at the species level to understand the impact of environmental factors, but intraspecific diversity has largely been ignored. To address this significant knowledge gap, we took advantage of the simple, culturable microbiome of *Drosophila*. First, we documented that natural populations of *D. simulans* harbor three diverged clades of *Lactiplantibacillus plantarum*, a key nutritional symbiont. We studied the distinct ecological roles of these three clades by exposing flies with their native microbiome to two temperature regimes in the laboratory. Tracking the three clades within the complete *Drosophila* microbiome over a period of more than 10 years at two temperatures, we identified strikingly distinct dynamics in response to the selection regime. We confirmed the functional differentiation of the three clades using *in vitro* growth measurements and *in vivo* mono-association assays. Our results highlight that environmental selection operates at the subspecies level. Therefore, we conclude that the functional diversification of the microbiome can only be understood when intra- and interspecific diversity is considered.

## Introduction

Intraspecific diversity has largely been overlooked in microbial ecology surveys. This can be in part attributed to the success of 16S rRNA profiling, which allows the characterization of species diversity with a very high efficiency (1,2), but cannot identify intraspecific diversity (3). Although environmental variation has been identified as one of the major factors driving species composition, its role for intraspecific adaptation has been largely overlooked (3). This blind spot of microbiome research is particularly surprising as it is known that conspecific strains, i.e. strains from the same species, can display large phenotypic variation (4). Clinically relevant species like *Escherichia coli*, in which different subspecies can be either pathogenic or commensal are a particularly illustrative example (5).

Although a few studies in environmental microbial communities have provided evidence for the functional importance of intraspecific variation (6,7), such analyses are considerably more difficult. Therefore, descriptive correlative studies are more common than functional assays (8). In the gut-dwelling bacterium *B. fragilis* conspecific strains compete for the same niche leading to the mutual exclusion of toxicogenic and non-toxicogenic strains in the gut environment (9). Not only in the human gut, but also in lakes, soil, and the sea conspecific strains frequently co-occur in sympatry (10–12). It has even been even reported that strain richness, i.e. sympatric strain diversity within the same species, stabilizes microbial communities in fluctuating environments (13,14).

In this work, we further explored the functional implications of intraspecific diversity within the same microbial population. We focused on the species *Lactiplantibacillus plantarum*, an extremely versatile lactic acid bacterium that has been associated with a variety of different habitats, including plants, the gastro-intestinal tracts of mammals and insects, as well as food such as meat, dairy, and pickled products (15). *L. plantarum* is a facultative symbiont of *Drosophila* (16–19). *Drosophila* larvae can enhance the growth of *L. plantarum,* which in response releases essential nutrients for the larvae under poor diet conditions (18,20). As a result of this nutritional symbiosis, *L. plantarum* is a prevalent taxon in the *Drosophila* microbiome (17).

To study the implications of intraspecific diversity of *L. plantarum* in fruit flies, we took advantage of an experimentally evolved *Drosophila simulans* population that has been adapting to two novel temperature regimes, hot and cold, for more than ten years. We detected three clades of *L. plantarum* co-occurring within the native microbiome of *D. simulans* population before the experimental temperature was modified. However, the exposure of the fly population to the novel temperature regimes altered the intraspecific composition of *L. plantarum,* revealing functional differences between the co-occurring clades. These functional differences were experimentally validated *in vitro* and *in vivo*. Our results provide a clear example for functional diversification of intraspecific diversity and demonstrate how experimental evolution can uncover this otherwise hidden functional diversity.

## Results

### Genome sequencing reveals the presence of three *L. plantarum* clades in the experimentally evolving populations

We sampled 33 isolates from the Florida experiment (See Materials and Methods for details) to characterize the diversity of *L. plantarum* (Figure 1, Table S1). The isolates formed three clades based on the pairwise average nucleotide identity (ANI). Between-group ANI values were 98.4-99.17%, below the ∼99.5% threshold for conspecific strain identity (21). On the other hand, the identity within groups was 99.48-99.99% (Figure 2). The clustering was consistent irrespective of whether we used SNPs in the core genome or presence/absence patterns of accessory genes. This indicates that the three clades are old with very limited genetic exchange.

**Figure 1.**
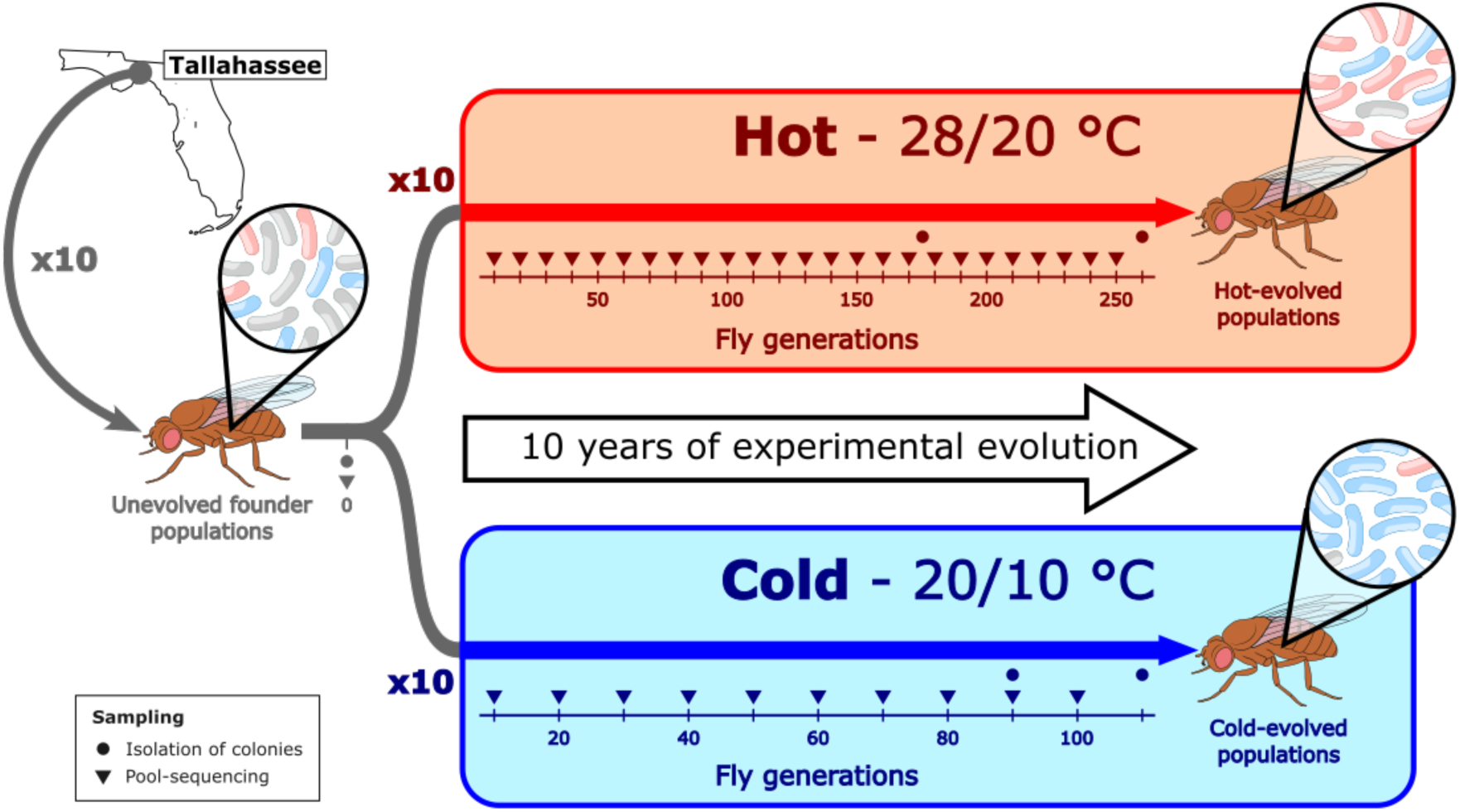
Experimental setup to study the long-term evolution of *L. plantarum* in *Drosophila simulans* populations.

**Figure 2.**
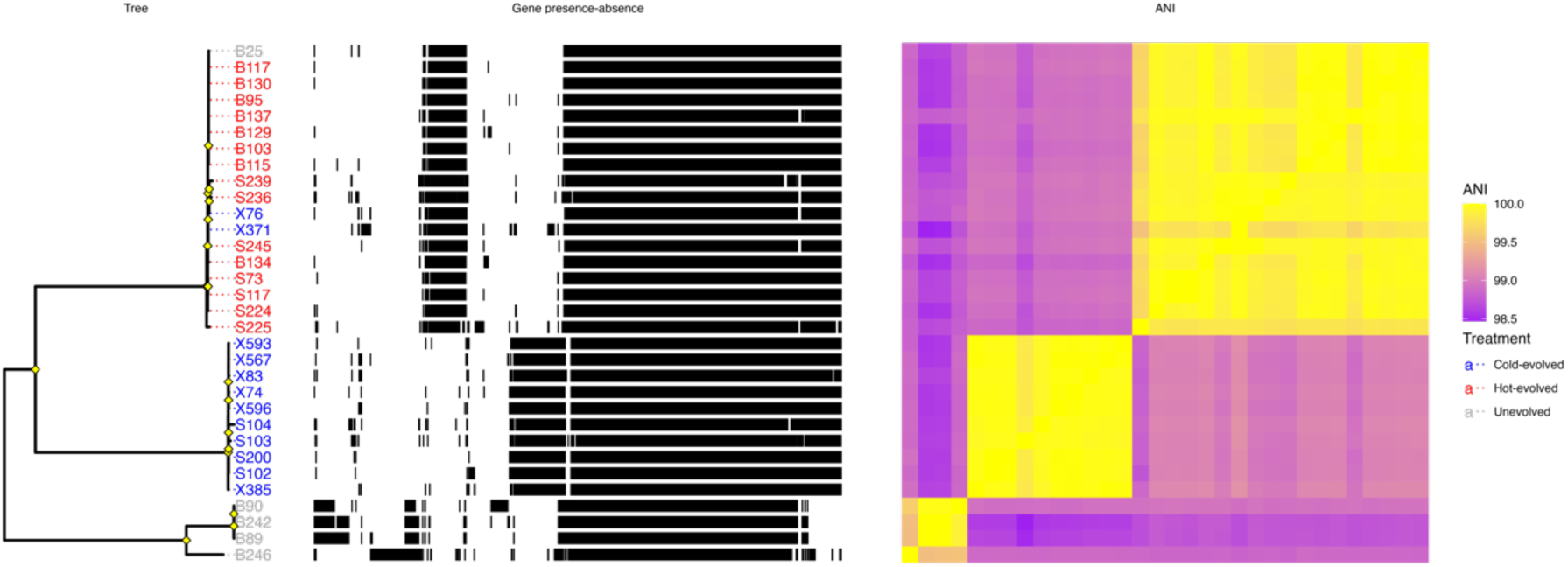
Pangenome of *L. plantarum* genomes isolated from the Florida experiment (Table S1), sorted by phylogeny. Left panel: Maximum likelihood tree built based on the core genome alignment. Tree leaves correspond to the name of the isolate and are coloured based on isolation origin: ancestral population (grey), hot-evolved population (red) or cold-evolved population (blue). Nodes supported by bootstrap values ≥ 90% based on 1000 replicates are highlighted in yellow. Middle panel: Pattern of genes presence (black)/absence (white) in each genome. Right panel: pairwise average nucleotide identity between the isolates. The colour ranges between 98.5% (purple) and 100% (yellow).

Since the isolates were sampled either from the unevolved, hot-evolved or cold-evolved populations (Figure 1), we tested whether the clades were randomly distributed across the experimental treatments and observed a non-random distribution (Fisher’s exact test, 3x3 contingency table, p = 5.5e-10). In combination with the clustering of clades (Figure 2), we conclude that each of the three clades is more prevalent either in the novel experimental populations or in the unevolved founder population, at the beginning of the experiment. We therefore coined the terms H (hot-evolved), C (cold-evolved), and U (unevolved) clades, that will be used throughout the manuscript.

In order to further investigate the association between *L. plantarum* genotype and temperature, we sampled from two additional experiments, obtaining seven isolates from hot-evolved *Drosophila simulans* Portugal and two isolates from constant-cold-evolved *Drosophila simulans* South Africa (Table S1). In a phylogenetic tree, the hot-evolved isolates and the constant-cold-evolved isolates grouped into clades H and C, respectively (Figure S1). These results do not only support the association between *L. plantarum* clade composition and environmental temperature (Fisher’s exact test, 3x3 contingency table, p = 2.6e-12), but also suggest that clade C is favored irrespective of whether the temperature is fluctuating (20 °C during the day, 10 °C at night) or constant (15 °C). Furthermore, the grouping of isolates originating from different experiments to the same clades indicates that these are not restricted to a single natural *D. simulans* population, but represent global diversity.

#### The three sympatric clades independently adapted to their host

Our analyses suggest that all three clades are shared among natural *D. simulans* populations, but it is not clear whether they diverged in *D. simulans* or independently colonized their *Drosophila* host. We used 79 publicly available *L. plantarum* genomes from several environments and geographic locations (Table S2) to reconstruct the phylogeny based on the core genome.

All three *L. plantarum* clades are more closely-related to genomes from other sources than to each other, which implies that they diverged before colonizing *D. simulans* (Figure 3). Moreover, the genomes from the clade H are almost identical to public sequences isolated from global collections of *Drosophila melanogaster* (Figure 3, Table S2). The clade U isolate B246 is closely related to the *Drosophila*-associated isolate LpWF originating from a wild *D. melanogaster* individual (22). Clade C did not have a close relative in the collection. The high similarity of the *L. plantarum* sequences isolated from *D. melanogaster* and *D. simulans* highlights that both *Drosophila* species share this component of the microbiome. Most likely a deeper sampling of the *D. melanogaster* microbiome will also detect clade C.

**Figure 3.**
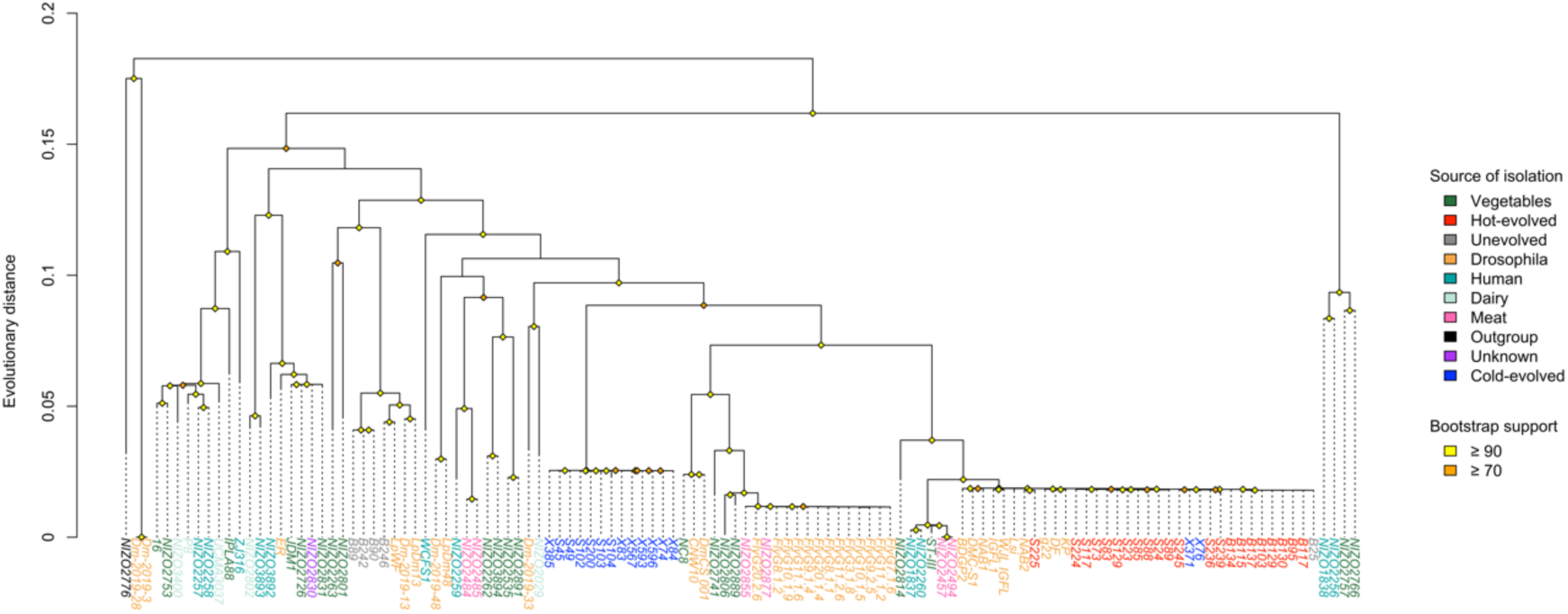
Maximum likelihood tree of 92 *L. plantarum* genomes. Tree leaves are coloured by source of isolation. The genomes from our lab are coloured according to experimental treatment: unevolved in grey, hot-evolved in red, and cold-evolved in blue. Nodes supported by bootstrap values based on 1000 replicates are highlighted in orange (≥ 70% bootstrap support) or yellow (≥ 90% bootstrap support).

Despite being phylogenetically divergent, the three sympatric clades could have converged functionally in the process of adaptation to a common environment. In order to test this, we built the *L. plantarum* pangenome including all the available isolates and performed hierarchical clustering based on the presence/absence of accessory genes. We found no grouping of the three sympatric clades (Figure S2), which suggests that they are not only phylogenetically diverged, but have the potential to be functionally different based on their pool of accessory genes.

#### Clade dynamics are temperature-specific and consistent between population replicates

Up to now, our analyses were restricted to isolates sampled at specific time points of the experiment. Since genomic DNA from flies with their microbiome was sequenced throughout the entire experiment (Figure 1), a metagenomic analysis could track the frequency trajectories of the three clades to shed more light on the adaptation process. We calculated the proportion of reads assigned to each clade in the population at a given time point. At the beginning of the experiment, the populations were dominated by clade U (54.5% mean relative abundance), followed by clade C (31.9%), and then clade H (13.6%). When the fly populations were subjected to the experimental treatments, clade H became dominant under the hot regime and clade C became dominant under the cold regime (Figure 4). Clade U greatly decreased in relative abundance in both regimes. This suggests that changes in environmental temperature alter the fitness landscape of the population and lead to a new adaptive state, in which the relative fitness of the clades change.

**Figure 4.**
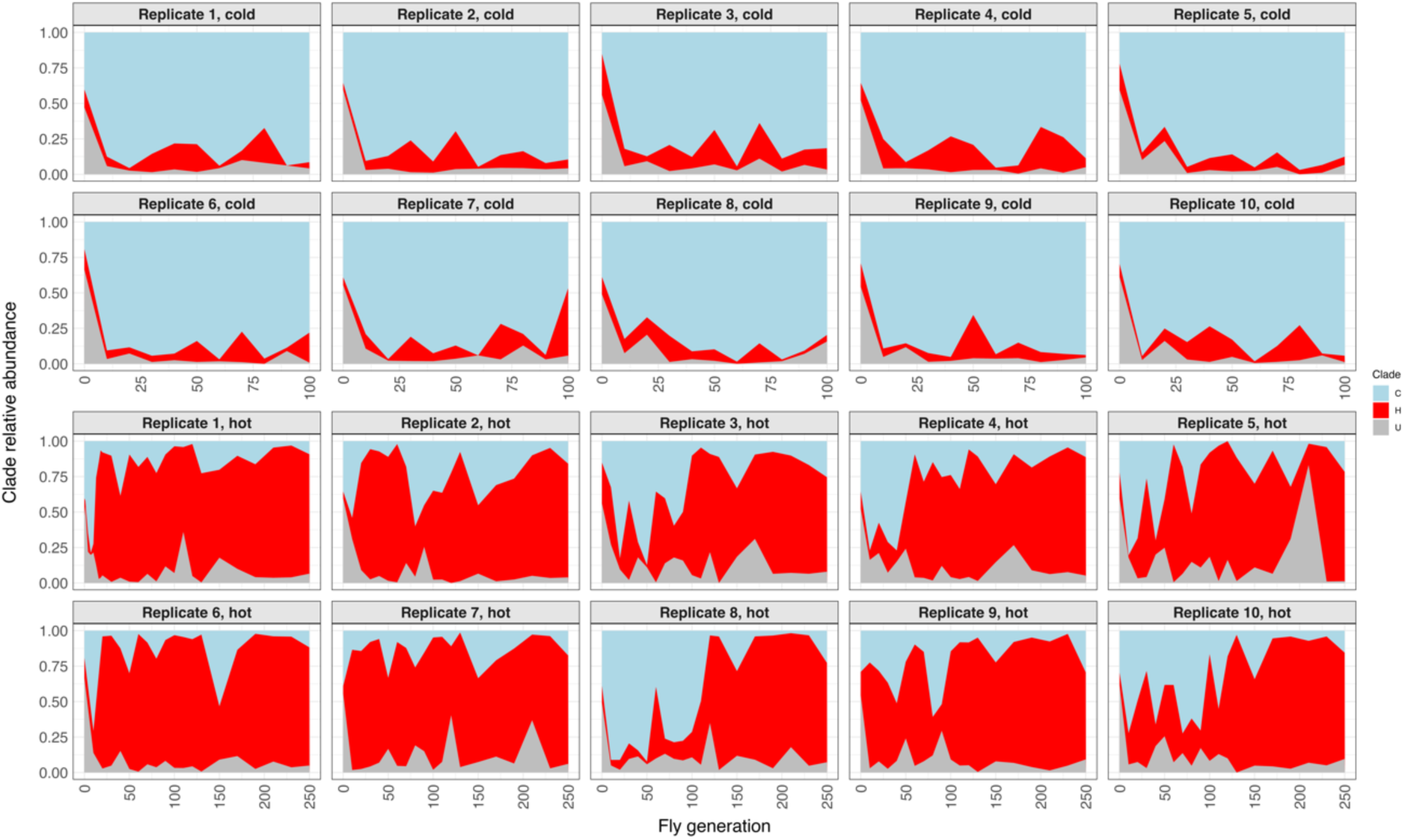
Clade composition over time in each replicate population from both temperature regimes. C in light blue, clade H in red and clade U in grey. Clade relative abundance was inferred by mapping competitively short reads against the three clades’ reference sequences.

A strong asset of the experimental evolution set up is the availability of 10 replicate populations that were generated from the same founder population and independently maintained throughout the entire experiment under the same conditions. Hence, the comparison of replicates provides an estimate for the strength of deterministic and stochastic forces during the experiment. In the cold regime, all ten replicates exhibited a rapid takeover of clade C (Figure 4). In contrast, the hot-evolved replicates were more variable in the rate at which clade H increased in abundance. The most extreme cases were replicates 8 and 10, in which clade C dominated the population for 90 generations before being replaced by clade H. Thus, the populations subjected to the hot regime took longer to reach a new equilibrium, which is also less reproducible than that of the cold regime. Overall, the temperature-specific dynamic is consistent across replicates in both regimes, confirming the association between environmental temperature and clade frequency that was observed during the isolation of the clones (Figures 1, 2 and S1).

#### Clades differ significantly in their growth kinetics in liquid culture

The metagenomic analysis showed that a different clade dominated each novel temperature regime and this was highly consistent across replicates. The temperature-specific dynamics can arise from clade-specific growth kinetics, as observed for *B. cereus* groups (23). Alternatively, temperature could affect *L. plantarum* indirectly, by acting on the host, the food, or other microbiome members. To determine if the observed clade dynamics were driven solely by differences in optimal growth temperature, we grew isolates from the three clades in liquid MRS medium at temperature conditions similar to the experimental settings (See Materials and Methods for details). The selected isolates represented all sampling events across the two temperature regimes (hot and cold) as well as the unevolved population (Table S1).

The three clades differed significantly in growth rate, carrying capacity, and inflection time, i.e., the time it takes an isolate to reach the log phase, in both temperature conditions (Figures 5 and S4; Kruskal-Wallis test, p < 0.005 for all parameters). However, the performance of the three clades did not fully match our expectations, based on the time series. In hot conditions, clade U outperformed clade C in growth rate, carrying capacity, and inflection time (Figure 5) despite clade C prevailed in the *Drosophila* populations for many fly generations (Figure 4). In cold conditions, clade C had significantly lower growth rate than clade H and no significant differences in carrying capacity and inflection time. These results reveal differences in growth kinetics of the three clades, but do not support the hypothesis that differences in optimal growth temperature caused the shifts in composition observed in the time-series experiments in flies.

**Figure 5.**
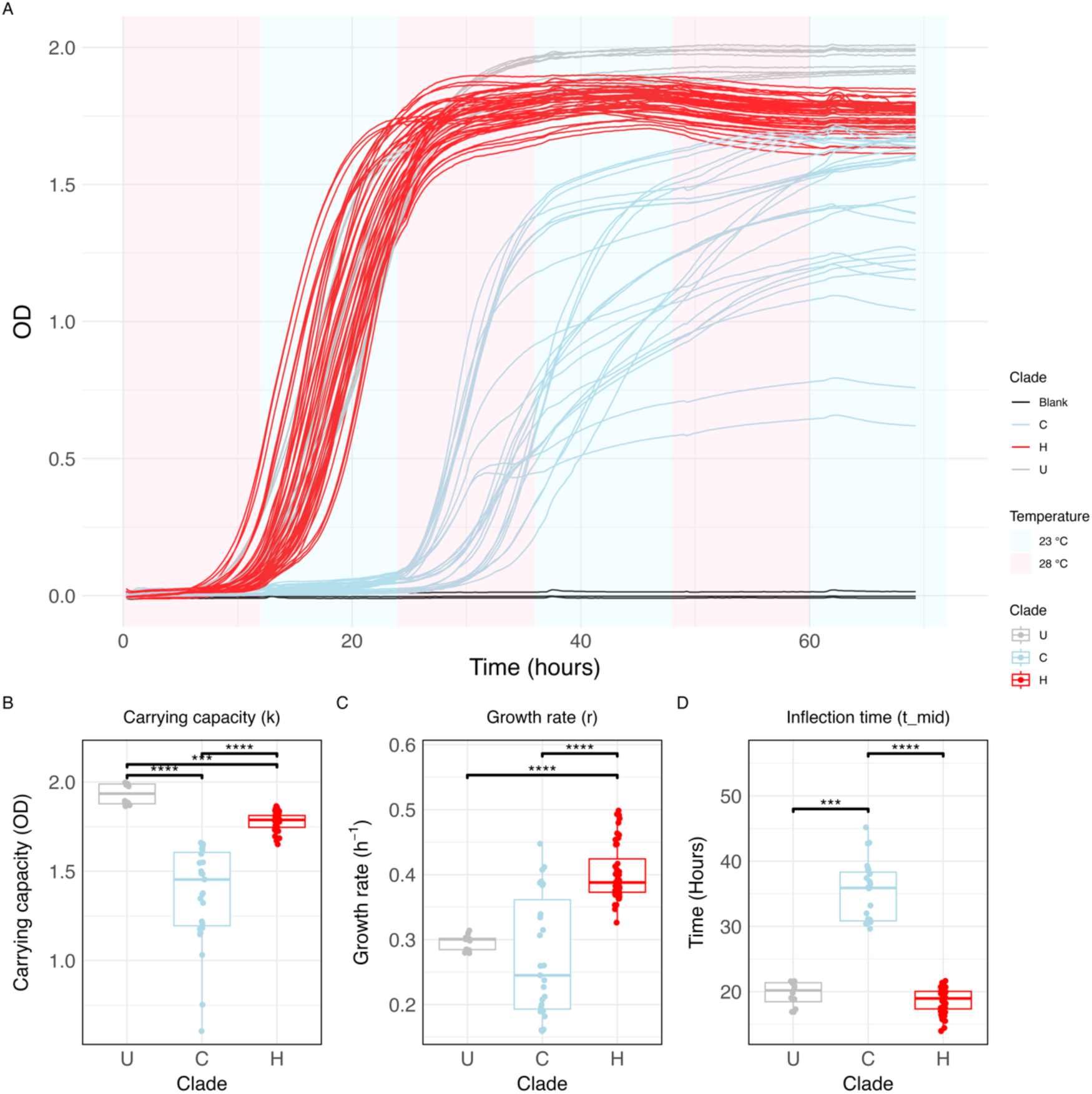
Clade-specific growth dynamics at hot fluctuating 28/23 °C. (A) Overlapped curves of all replicates. Lines are coloured by clade. Background colour represents the growth temperature at a given time. (B-D) Boxplots depicting the carrying capacity, growth rate, and inflection time of each clade. We measured the growth of four isolates from clade U, nine isolates from clade C and sixteen isolates from clade H (Supplementary table S1). Each isolate was grown three time. Each dot corresponds to a technical replicate. Data is grouped and coloured by clade. Statistical significance was determined using Dunn’s test with Holm-adjusted p-values. Only significant comparisons are indicated. **** *p* < 0.0001; *** *p* < 0.001; ** *p* < 0.01; * *p* < 0.05

Nevertheless, we caution that the environment in the time-series experiment is radically different to that of liquid MRS (24). We confirmed this by agar-drop assays in solid fly food, which resulted in delayed growth of clade U, relative to clades H and C, in both focal temperature regimes (Figure S12). Furthermore, *L. plantarum* lives in a complex community in which the different clades interact with each other, with other members of the microbiome, and finally also with the host. Bacteria can persist in the community by thriving in the food or by stably colonizing the fly gut. The former strategy might be controlled by growth rate and carrying capacity, but the latter is affected by additional factors such as death rate in the gut and defecation rate, that cannot be assessed in liquid medium (22). Thus, observed differences in growth could indicate different strategies to persist in the community.

### Fitness effects of *L. plantarum* strains on their host

Although growth dynamics differ significantly among the three clades in growth culture, this does not explain the observed changes in abundance during experimental evolution. Therefore, we turned to the *Drosophila* host to explore the fitness consequences of colonization by each clade.

*L. plantarum* can increase larval fitness of *Drosophila melanogaster* relative to germ-free flies under low protein conditions (18,20,25). Protein content is controlled by the amount of yeast in the fly food (18,20,25). *L. plantarum*-mediated benefits have been observed in low yeast (<=12 g/l) food, but not in protein-rich diets (50-80 g/l yeast). Because our experimental diet contains an intermediate yeast content (24.3 g/l), we wondered whether we could detect temperature-dependent fitness effects of the host in this food. To test this, we designed a set of inoculation experiments in both focal temperatures using two host species, *D. melanogaster* and *D. simulans*, and the three *L. plantarum* clades. While we could produce axenic *D. melanogaster*, attempts to produce axenic *D. simulans* were unsuccessful. Thus, *D. simulans* used in these experiments were dechorionated but not axenic.

Contrary to experiments with protein-poor diets, none of the *L. plantarum* clades provided a fitness advantage to the host relative to controls in any of the species. In the case of clades U and H, the number of offspring and developmental time did not differ significantly from the dechorionated controls (Figure 6; Dunn’s test, p > 0.05 for all pairs). Surprisingly, in both host species we observed significant fitness reductions in the presence of clade C. While *in D. melanogaster* offspring number decreased and developmental time increased at both temperatures (Figure 6; Dunn’s test, p < 0.05 for all significant comparisons), in *D. simulans* only developmental time increased in the cold only (Figure 6; Dunn’s test, p < 0.05 for all significant comparisons). We attribute the weaker effects in the non-axenic flies to a reduced *L.* plantarum load due to the presence of other taxa (22), or to higher-order interactions with other members of the microbiome (26–28).

**Figure 6.**
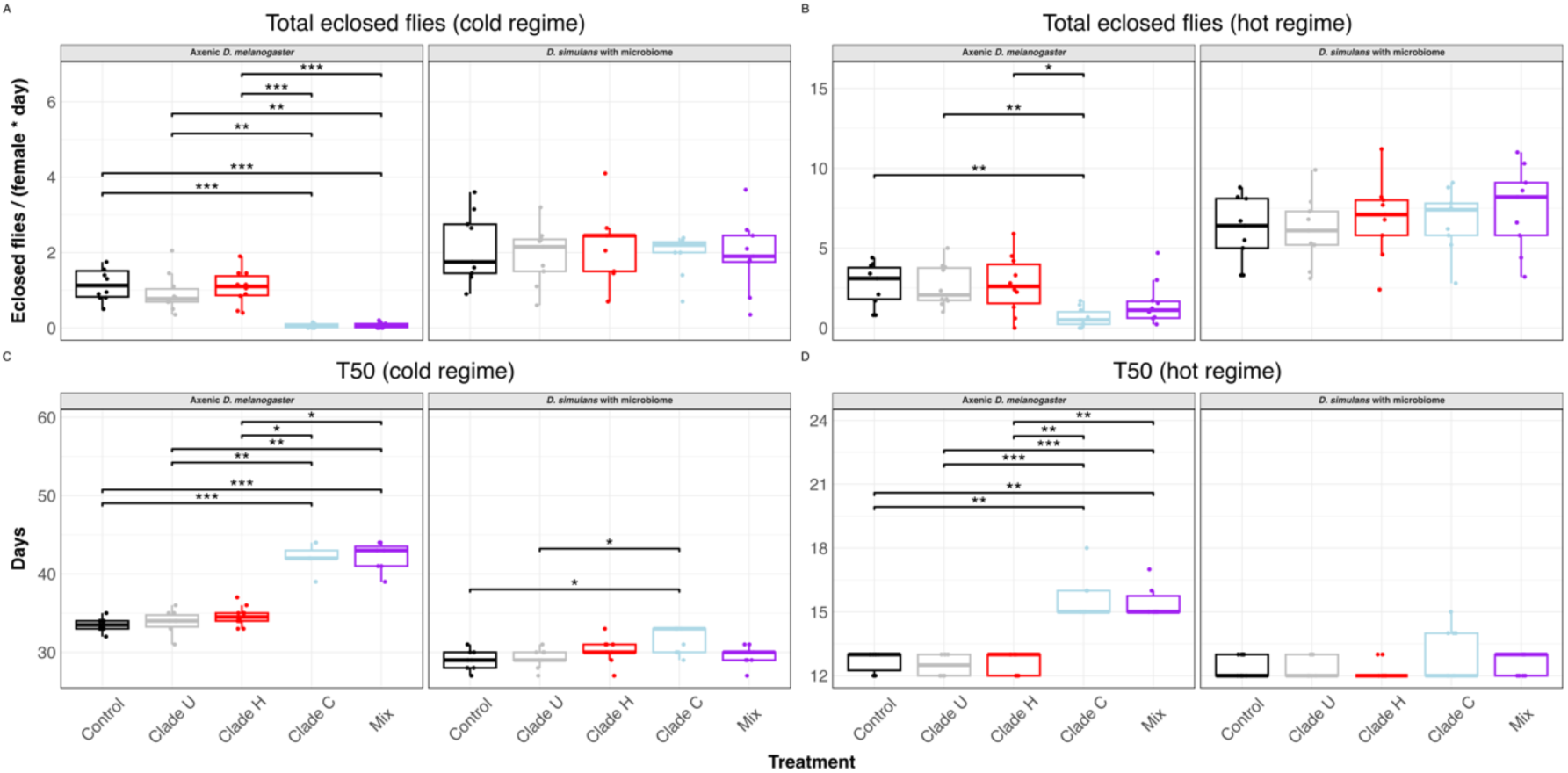
Fitness effect of *L. plantarum* inoculation in axenic *D. melanogaster* and conventionally reared *D. simulans* in the first transfer (See Materials and methods). (A, B) Total number of F1 flies eclosed normalized by day and female under the cold (A) and hot (B) regime. (C, D) Developmental time, estimated as the number of days it takes 50% of the offspring to eclose. Measurements were grouped by inoculation treatment and *Drosophila* species. Each dot corresponds to a biological replicate (n = 10 for *D. melanogaster* and n = 9 for *D. simulans*). Statistical significance was determined using Dunn’s test with Holm-adjusted p-values. Only significant comparisons are indicated. *** p < 0.001; ** p < 0.01; * p < 0.05.

We further explored the role of bacterial load on the host fitness by extending the experiment to a second transfer: the same parents laid eggs into a new set of sterile vials (See Materials and Methods for details). The effect of clade C on the host disappeared for *D. simulans* and was diminished in *D. melanogaster* relative to the first transfer (Figure S5; Dunn’s test, p < 0.05 for all significant comparisons). Since the bacterial load could be a major factor influencing these results, we performed an additional experiment in *D. simulans* to quantify the bacterial load of parental flies in the second transfer. Median bacterial load of flies inoculated with clade C was 4.1×10^5^ colony forming units (CFUs) per fly in the hot regime and 1.3×10^4^ CFUs/fly in the cold one, at least 10-fold higher relative to the other clades in the same regime (Figure S6). These results suggest that effect of clade C on the host depends on the bacterial load, and could be related to an excess of bacterial growth inside the host.

In summary, *L. plantarum* does not confer a fitness benefit to the host in any temperature regime. Moreover, clade C, which is dominant in the cold-evolved populations (Figure 4), is clearly detrimental for reproduction and development in *D. melanogaster* hosts. In *D. simulans*, though, slower development provided by clade C could be beneficial in the cold, as observed in other environmental conditions (29,30).

#### Functional divergence on the genomic level

Our analyses clearly indicate functional divergence between the three clades at genetic and phenotypic levels (Figures S2, 5 and 6). Therefore, we were interested in identifying the genes that contribute to this functional differentiation. A comparative KEGG pathway analysis showed that clade C contains the most KEGG Orthologs (KOs), 1163 KOs, followed by clade H (1157 KOs), and clade U (1131 KOs). A subset of 111 orthologs is unique to one or two clades. Of these, 41 are shared by clades H and C (Figure S7). These differential orthologs fell into 53 KEGG metabolic pathways.

Interestingly, clades C and H encode a shared genetic repertoire related to sugar metabolism that is absent in clade U, which does also not encode any private sugar-related function (Table S3). Clade U cannot use sorbitol as carbon source, since it does not encode the PTS transporter to internalize it (EC: 2.7.1.198; Figure S8) nor the enzyme sorbitol dehydrogenase (EC: 1.1.1.140), that converts sorbitol into fructose (31). Clade U also lacks transaldolase activity (EC: 2.2.1.2), that connects the Embden–Meyerhof–Parnass pathway with the non-oxidative pentose phosphate pathway (32). Finally, clade U is unable to hydrolyze chitobiose into N-acetyl-glucosamine monomers because it lacks the enzyme hexosaminidase (EC: 3.2.1.52).

Additionally, clade H and C harbor different sugar-related functions (Table S3), indicating that, despite partial overlap, these strains possess distinct sugar metabolic capacities. We also identified functional differences between clades in cofactor biosynthesis and respiratory metabolism. These differences do not explain the clade-specific selection, but reflect the different evolutionary histories of the clades (Table S3, figures S8-S11). A detailed description of the functional differences between clades can be found in the supplement.

To sum up, among the clade-specific genes we found an enrichment of sugar-related functions in clades C and H. This, summed to the delayed growth of clade U in solid fly food (Figure S12), suggests that diet, in addition to temperature, affected the intraspecific dynamics of *L. plantarum*.

## Discussion

Intraspecific diversity within the same environment has been reported to be ecologically important to maintain the stability of the entire microbial community (11,13,14). This parameter has been studied in lakes, soils, seas, and the human gut (10–12). However, little attention has been given to intraspecific variation in *Drosophila*, and insect microbiomes in general (8). Based on our results in experimentally evolved fruit flies, we propose that within-species competition, thus far largely overlooked, could contribute to ecological adaptation and evolution of the host, as it has been observed at the species level in *Drosophila* (33) and other animals (34–36).

Our finding of three co-occurring subspecies of *L. plantarum* in our populations is particularly surprising, given the low microbiome richness of *Drosophila* at species level, which is estimated to be orders of magnitude lower than in humans (37,38). However, the intraspecific richness of *L. plantarum* in our flies was three times higher than that estimated in human gut microbiomes (11). We note that a recent study also found several co-occurring *L. plantarum* variants within a *Drosophila* population (39). Our work further expands this observation by demonstrating that this diversity is not neutral, but rather driven by external factors. This discrepancy between inter- and intra-species diversity suggests that the *Drosophila* microbiome is more complex than previously thought (16,26,40). However, this diversity exists below the species level and has therefore remained cryptic due to the limitations of amplicon sequencing (3).

By altering the experimental temperature at which the populations were maintained, we discovered consistent shifts in the composition of *L. plantarum* subspecies, that inevitably led to the dominance of a single clade. Further *in vivo* experiments revealed that clade composition significantly impacts the fitness of the host. It was previously shown that nutritional symbiosis in *L. plantarum* is strain-specific (19), and that some genotypes can be detrimental for the host (41,42). In our study, we also showed that clade C was detrimental to the host, but nevertheless in the *Drosophila* time series experiment, this clade outcompeted the other clades, which did not have a negative effect on host fitness. This demonstrates that the benefits of parasitism or mutualism in the *Drosophila* microbiome are context-dependent, as already observed in humans (43,44).

Finally, we also gained insight into how the microbiome of the fly could have adapted to laboratory conditions. The two clades, C and H, which are favored in the time-series, harbor unique sugar-related metabolic functions, consistent with faster growth in the sugar-rich laboratory diet. Besides the novel temperature regimes, introduction to the laboratory changes the diet and mobility of the host and the microbiome relative to the wild (45,46). This explains the rapid decrease of clade U in both temperature regimes, despite showing high fitness in liquid MRS medium. Our findings are consistent with microbiome laboratory adaptation, and agree with previous observations in *L. plantarum* (17) and *Acetobacter* (46).

Overall, our work emphasizes the importance of intraspecific diversity in maintaining the stability of microbial communities and the host’s ability to adapt to environmental changes. The current climate crisis has sparked a significant interest in how environmental temperature affects microbial communities at the species level (47–49). Our study highlights that even subspecies diversity plays a key role in adaptation to environmental temperature, and probably other abiotic factors. Therefore, we propose that a comprehensive understanding of adaptive processes and ecological dynamics in the case of host-microbiome interactions depends critically on including diversity at the subspecies level.

## Materials and Methods

### Experimental evolution

Twenty replicate populations were set up using 202 isofemale lines from a natural *D. simulans* population collected in Florida (50). The populations were maintained in a 12 h photoperiod and at two different novel temperature regimes: ten replicates were kept in a fluctuating hot environment (28 °C during the day and 18 °C at night), whereas the other ten were kept in a fluctuating cold environment (20 °C during the day and 10 °C at night). The census population size of the replicates was 1000-1250 with a 50:50 sex ratio. The flies in each replicate were equally distributed across five 300 ml bottles containing 60 ml of standard *Drosophila* medium (300 g Agar + 990 g sugar beet syrup + 1000 g malt syrup + 2,310 g corn flour + 390 g soy flour + 900 g yeast in 37.5 L water) (51). The evolved *D. simulans* populations were sequenced in 10 generation intervals using Pool-Seq (52,53). A subset of these sequencing reads has been used to study the evolutionary dynamics of fruit flies (53). Similar pool-seq time series data have been previously used to analyze long-term endosymbiont and microbiome dynamics in *D. melanogaster* populations evolving in the same experimental conditions (54–56).

We also isolated *L. plantarum* from two other experimental evolution experiments, one originated from a South African population reared at constant 15 °C and the other from Portugal, which was reared in the fluctuating hot regime as described earlier (57). Flies in both experimental evolution studies followed the same maintenance regime as the Florida populations, with temperature being the only experimentally altered variable (58).

#### Bacterial isolation

We primarily obtained data from the experimental evolution studies using the Florida *D. simulans* founder population and collected 33 *L. plantarum* genomes from five different time points; two time points from each experimental temperature and one time point corresponding to the ancestral flies (Table S1). In addition, we obtained two and seven genomes from experimental evolution studies originating from South African and Portuguese *D. simulans* populations.

Around 100 flies from each population were homogenized in sterile PBS using an autoclaved pestle. The homogenate was diluted to avoid physical contact between colonies and streaked on agar plates with three different media: De Man, Rogosa, and Sharpe (MRS), mannitol and tryptic soy. The plates were incubated at 28 °C for 48 h. We identified *L. plantarum* using PCR with custom *Lactiplantibacillus*-specific primers (in 5’-3’ orientation, LacF: GATGGGCGCTTACCCGATTA, LacR: CTGCCCCGCAAATTGTTTCA). Colonies which could successfully be amplified were picked and regrown to stationary phase in the same medium from which they were isolated, but in liquid format. A glycerol stock of each isolate was made by mixing the culture with glycerol (50% v/v) in 1:1 proportion. The proportion of *L. plantarum* colonies, relative to those of other taxa, varied across growth media and fly populations. A list of other microbial taxa isolated from the flies can be found in Table S4.

#### DNA extraction and sequencing

Genomic DNA was extracted from all the replicate populations using the high salt method (59). The fly populations were sampled in 10 generations intervals starting from generation 0. At sampling the age of the flies varied between four and eight days for the hot environment and between nine and 16 days for the cold environment. Pools of flies were sequenced at various time points, using a range of library kits, insert sizes, and read lengths (Table S5). We expanded the genomic dataset used in Barghi *et al.* 2019 (53) with a new set of pools covering the hot-evolved populations up to generation 250 and the cold-evolved population until generation 100.

For sequencing, *the L. plantarum* isolates were grown to stationary phase in MRS medium. gDNA was isolated with the high salt extraction method (59). Additionally, a lysozyme pre-treatment was used to degrade Gram-positive cell wall (60). Briefly, the pellet was resuspended in 480 µl of EDTA 50 mM. The suspension was treated with 120 µl of lysozyme (20 mg/ml dissolved in NaCl 30 mM – EDTA 2mM) and incubated for two hours at 37 °C. After this, the samples were centrifuged at maximum speed, the supernatant was discarded, and the pellet was further treated following the high salt gDNA extraction. Genomic DNA was quality-controlled on an Agilent Bioanalyzer (Agilent Technologies, Inc., Santa Clara, CA) and subsequently used to prepare DNBSEQ short-read libraries, and 2x150bp reads were sequenced on a DNBSEQ-G400 (MGI Tech Co., Ltd., Shenzhen, China).

#### Genomic assemblies

Sequencing reads were processed using BBDuk (*BBMap*, 2025 v38.90) to remove adapters, ΦX174 sequences and low-quality bases with the following parameters: ktrim=r, k=23, mink=11, hdist=1, tbo, tpe. Reads were assembled using SPAdes v4 (62) with kmer sizes 21, 33, 55, 77, 99, and 127. Subsequently, CheckM2 v1.0.1 (63) was used to estimate the completeness and contamination of each assembly, and the taxonomy was determined with GTDB-Tk v2.1.1 (64). Ten assemblies had more than 5% contamination, probably due to accidentally picking two adjacent colonies. We used Centrifuge v1.0.4 (65) to classify and exclude contigs with exogenous origin from these assemblies.

#### Comparative genomics and phylogeny

We built the pangenome and phylogenetic tree of the collection of genomes isolated from our lab (Table S1). First, we used Prokka v1.14.6 (66) to predict the genes in all the genomes. Roary v3.13 (67) was used to construct the isolates’ pangenome and extract the aligned genes belonging to the core genome, i.e. those that are present in all genomes. SNP-sites v2.5.1 (68) was used to extract the variable positions from the concatenated core genes. A maximum likelihood tree of the aforementioned variable positions was built using the model GTR+ASC and 1000 bootstraps with IQ-TREE v3.0.1 (69). Additionally, pairwise average nucleotide identity between all the isolates was computed using the program PyANI v0.2.13.1 (70), with ANIb as alignment method using the whole genomes. This workflow was carried out for the subset of 33 genomes isolated from the Florida experiment and for the whole dataset of 42 genomes, including those isolated from South Africa and Portugal experiments. Unless specified otherwise, subsequent analyses were conducted using the whole dataset pangenome.

The same phylogenetic and pangenomic methods were used for the extended *L. plantarum* genomes collection, including 79 publicly available genomes (Table S2). In order to root the tree, an additional tree was made including the reference genome from the sister species *Lactiplantibacillus paraplantarum* (GCA_003641145.1) as outgroup. The *L. plantarum* isolate that was most closely related to *L. paraplantarum*, NIZO2776, was used to root the tree.

In order to investigate which metabolic functions were differentially encoded in each clade, we followed the Anvi’o v7.1 pangenomic workflow (71,72) to identify “gene clusters” that are unique or shared across the genomes isolated in our lab. These gene clusters were further annotated against the KEGG database using the built-in Anvi’o function ‘anvi-run-kegg-kofams’. The script ‘anvi-compute-functional-enrichment’ (73) was then used to screen for KEGG functions that are differentially enriched between the clades. Briefly, this script estimates the fraction of genomes from each clade that encode each KEGG Ortholog (KO) and tests whether they are significantly associated to one or more groups. We used the R package ggKegg v1.4.1 (74) to visualize the KEGG pathways in which at least one gene was enriched.

#### Competitive mapping and clade relative abundance calculation

First, we pre-filtered the *Drosophila* pool-seq reads by removing the reads that have eukaryotic or *Wolbachia* origin. We used Bowtie v2.5.4 (75) with stringent settings (-D 500 -R 40 -N 0 -L 20 -I S,1,0.50 --no-mixed --no-discordant) to map each pool-seq data set against a collection of reference genomes that included *D. simulans* (GCF_016746395.2), *D. melanogaster* (GCF_000001215.4), *D. mauritiana* (GCF_004382145.1), *H. sapiens* (GRCh38), *M. musculus* (GCF_000001635.27), *A. thaliana* (GCF_000001735.4), *S. cerevisiae* (GCF_000146045.2), *C. lupus* (GCF_011100685.1) and a several insect-infecting *Wolbachia* strains (GCF_000008025.1, GCF_000022285.1, GCF_000376585.1, GCF_000376605.1, GCF_000475015.1). We kept the read pairs in which none of the reads mapped to any of the genomes in the collection, which should predominantly represent prokaryotic sequences.

Then, we used SuperPang v0.9.6 (76) to build the reference graph pangenome of each *L. plantarum* clade using all the genomes belonging to the clade as input (Table S1, Figure 2). In order to calculate the clades relative abundance in each pre-filtered pool-seq, we used BBMap (*BBMap*, 2025 v38.90) with “perfect mode” settings to competitively map each read against the three concatenated clade pangenomes. For each sample, the number of reads that mapped uniquely to each clade was counted and normalized to Reads Per Million (RPM) in order to account for differences in genome size and sample depth. *L. plantarum* represented between 0.0438% and 17.2246% of relative abundance of the reads in all time points.

We benchmarked our approach to confirm the relative abundance estimates from our competitive mapping pipeline. Briefly, we simulated 200k HiSeq reads from each reference graph pangenome using InSilicoSeq v2.0.1 (77), and used ‘seqkit sampl’ v2.3.0 (78) to mix them in different ratios, maintaining the original size of 200k reads: 1:0:0, 0:1:0, 0:0:1, 1:1:1, 2:1:7, and 8:1:1. Each of the combined read sets was randomly subsampled from 150k to 10 reads (10, 20, 40, 60, 80, 100, 200, 300, 400, 500, 600, 700, 800, 900, 1k, 2k, 5k, 10k, 150k reads) and mapped competitively to the three clades. We confirmed that the relative abundances were correctly estimated with as few as 100-1000 reads mapping the references (Figure S3).

#### Growth assays

The glycerol stocks from all the available *L. plantarum* isolates were re-grown in agar MRS. A single colony per isolate was grown in liquid MRS and passaged daily for five days in order to remove the influence of freezing. On the fifth day, each suspension was normalized to an optical density (OD_600_) of 0.01, and serially diluted 1:100 in MRS. We selected 31 isolates that represented samples from all the available time points and temperature regimes, and plated them in triplicates in a flat-bottom 96-wells plate. Three wells were filled with sterile MRS medium as blank controls. Later, we detected contamination in two of the isolates, leaving us with 29 isolates. In total we grew four isolates from clade U, nine isolates from clade C and sixteen isolates from clade H (Supplementary table S1). The plate was incubated for 72 h and OD_600_ was measured every 15 minutes in a Synergy H1 plate reader. The temperature regime was chosen to be similar to the hot conditions of the fly populations, but due to limitations of the spectrophotometer’s cooling capacity instead of 28 °C during the day and 18 °C at night, we set it to 28 °C during the day and 23 °C at night. Orbital shaking was set to 425 cpm.

Growth in cold conditions could not be measured with the plate reader as it does not have cooling capacity and the experiment would have lasted more than a week, so we used an alternative approach. We grew twelve isolates, four from each clade (Table S1), in glass test tubes using 5 ml MRS. We normalized each suspension to an OD_600_ of 0.05 and then diluted 1:100 in 5 ml of MRS. We also prepared an MRS blank. We prepared six replicates per isolate and blank. Three replicates were set at 17:00 and three the next day at 10:00. All tubes were incubated at constant 20 °C and 168 rpm in a shaker (Excella E24, New Brunswick, USA). The OD of all tubes was measured for twelve days at 10:00 and at 17:00 using a spectrophotometer (Ultrospec 10; Biochrom, Cambridge, UK). By using the replicates at two different days, we effectively duplicated the measured time points without the need to increase the sampling frequency.

The R package GrowthCurver v0.3.1 (79) was used to infer the growth parameters from the growth curves. All statistical analyses were conducted in R v4.4.2 (80) using the package rstatix v0.7.2 (81). We used the Kruskal-Wallis test to assess overall differences in growth parameters among the clades. For post-hoc pairwise comparisons, Dunn’s test was applied, with p-value adjustments based on the Holm method. Differences were considered significant at p < 0.05.

#### Agar-drop assays

One representative isolate from each clade (Table S1, B89 for ancestral, S103 for cold, and S239 for hot) was grown overnight in MRS medium from glycerol stocks. OD_600_ was normalized to 0.2, and subsequently diluted 1:10 six times. We plated on fly food Petri dishes 10 µl drops of the original suspension and of 10^-2^, 10^-4^, and 10^-6^ dilutions. We incubated three plates at 28/18 °C and three at 20/10 °C for 10 days. Pictures of all plates were taken at incubation days 3, 6 and 10.

#### Production of axenic flies

We generated germ-free flies, using a modified version of the dechorionation protocol described by Kietz *et al.* (82). Specifically, we used 2.8% active chlorine bleach instead of 1% and supplemented bleach, EtOH, and deionized H_2_O with TritonX (1% v/v). The addition of this detergent prevents dechorionated eggs from sticking to the walls of the tubes, thus facilitating their transfer to bottles. The whole process was carried out in sterile conditions using a laminar flow hood. Axenic flies were maintained via periodic transfers to new sterile food inside the laminar flow hood. We first carried out this procedure with several *D. simulans* isofemale lines, but it was not possible to fully eliminate the residual microbiome despite several rounds of treatment. However, we could produce and maintain axenic *D. melanogaster* Oregon-R flies. For this reason, we decided to use *D. melanogaster* as axenic host in the inoculation experiments.

#### Inoculation experiments and measurement of bacterial load

The representative isolate from each clade (Table S1, B89 for ancestral, S103 for cold, and S239 for hot) was re-grown from the glycerol stocks in MRS medium. After three transfers, OD_600_ was normalized to 0.05. Autoclaved vials with axenic fly food were inoculated with 50 µl of each bacterial suspension. This volume corresponds to ∼2.5×10^5^ CFUs based on colony counting of serial dilutions. In total five treatments were used: the pure culture from each genotype, a mixture of the three genotypes in equal proportions (“mix”), and control, in which the vials were inoculated with 50 µl of fresh MRS medium. After overnight absorption of the liquid, twenty axenic flies (ten female and ten male flies) were added to each vial and allowed to lay eggs for either 24 h (hot conditions) or 48 h (cold conditions). After this step, the flies were transferred to a new set of vials with axenic, uninoculated food, and allowed to lay eggs for another 24 h (hot conditions) or 72 h (cold conditions). The adults were discarded and the vials were incubated at hot (28 °C during the day and 18 °C at night) or cold conditions (20 °C during the day and 10 °C at night) until the new generation eclosed. The number of eclosed flies was recorded daily. We estimated the total number of flies per vial and the developmental time as proxies of fitness. We calculated developmental time as the day from the start of the experiment until the day at which 50% of the flies in the vial were eclosed. The inoculation experiment was carried out with axenic *D. melanogaster* Oregon-R (ten replicates per treatment) and dechorionated, but non-axenic, *D. simulans* from Florida (nine replicates per treatment).

To quantify bacterial load of individual flies, we transferred four *D. simulans* males to two sets of inoculated vials as in the previous experiment. We allowed the flies to feed for 24 h in either hot or cold conditions before transfer to sterile food and maintained in the same regime for 72 h. Each fly was then placed in a 2 ml tube with 200 µl of PBS and a sterile 4 mm glass bead. The tubes were shaken at 1450 rpm for 4 minutes in a MiniG 1600 vertical shaker (SPEX®SamplePrep). Homogenates were diluted 1:10 (cold-treated) or 1:100 (hot-treated) in PBS, and 200 µl of each dilution was plated in an MRS plate. After overnight incubation at 28 °C, CFUs were counted in each plate. Based on colony morphology, most CFUs corresponded to *L. plantarum* in inoculated flies, despite using *D. simulans* harboring microbiome.

We used the Kruskal-Wallis test to assess overall differences among the treatments. For post-hoc pairwise comparisons, Dunn’s test was applied, with p-values were adjusted using the Holm method. Differences were considered significant at p < 0.05.

## Acknowledgments

We thank Cameron Strachan and Evelyne Selberherr from the Zentrum für Lebensmittelmikrobiologie (Veterinärmedizinische Universität Wien) for granting us access to a plate reader for the growth experiment. We also wish to thank Martin Polz, William Ludington, Cameron Strachan and Xiaoqian Annie Yu for their feedback regarding experimental design and interpretation of the results. We thank Rupert Mazzucco for his help with the processing and management of the genomic data from the lab. We thank Susamman Biswas for his help with the molecular characterization of the *L. plantarum* isolates. Finally, we are grateful to all members of the Institut für Populationsgenetik (Veterinärmedizinische Universität Wien) for helpful discussions and feedback throughout the project. Part of the Illumina sequencing was performed by the Next Generation Sequencing Facility at Vienna BioCenter Core Facilities (VBCF), member of the Vienna BioCenter (VBC), Austria (www.vbcf.ac.at).

## Author Contributions

Bosco Gracia-Alvira: Conceptualization, Data curation, Formal analysis, Investigation, Methodology, Resources, Software, Visualization, Writing – original draft. Stefanie Migotti: Resources, Writing – review and editing. Xiaomeng Tian: Resources, Writing – review and editing. Viola Nolte: Data curation, Resources. Christian Schlötterer: Conceptualization, Project administration, Funding acquisition, Resources, Supervision, Writing – review and editing.

## Competing Interest Statement

The authors declare no conflicts of interest.

## Funding

This work has been funded by the ERC (Achadapt) and the Austrian Science Fund (FWF, 10.55776/W1225, 10.55776/PAT7786824, 10.55776/F91).

## Supporting Information Text

### Extended clade-specific differences in KEGG metabolic pathways

Clades C and H encode a shared genetic repertoire related to sugar metabolism that is lacking in clade U. In contrast, the latter does not encode any unique sugar-related function. Clade U is unable to hydrolyze chitobiose into N-acetyl-glucosamine monomers because it lacks the enzyme hexosaminidase (EC: 3.2.1.52). It also lacks transaldolase activity (EC: 2.2.1.2), that connects the Embden–Meyerhof–Parnass pathway with the non-oxidative pentose phosphate pathway (1). Finally, it cannot use sorbitol as a carbon source, since it does not encode the PTS transporter to internalize it (EC: 2.7.1.198; Supplementary Figure S8) nor the enzyme sorbitol dehydrogenase (EC: 1.1.1.140), that converts sorbitol into fructose. These metabolic capacities could play a role in the rapid decrease in abundance of clade U observed in both temperature regimes (Supplementary Table S3). Chitobiose is the main product of chitin degradation, that makes up the flies’ exoskeleton (32). We have detected the presence of microbial chitinases in the metagenomic reads (unpublished observation), which suggests that chitobiose might be available in the community. Thus, we hypothesized that the ability to exploit this ubiquitous source of carbon and nitrogen could be very advantageous in the fly microbiome context (Figure 4), but would not affect the fitness in liquid MRS culture (Figures 5 and S4). Competitive advantage should be reflected in higher bacterial loads within individual flies, as bacteria in the gut are in contact with the peritrophic matrix (2). However, we found no significant differences between bacterial loads upon inoculation with clade H (harbours exosaminidase) and clade U (lacks hexosaminidase) (Figure S6). Thus, experimental data do not support that chitobiose consumption confers a competitive advantage to clades C and H. We also found other sugar-related functions are differentially present between clades C and H (Supplementary Table S3), suggesting that, despite having overlapping functions, these clades also encode unique sugar-related functions.

We also found differences in cofactor biosynthesis pathways. Although all the analyzed genomes can import riboflavin, the capacity to produce it *de novo* is enriched in clades C and H (Supplementary Table S3; Supplementary Figure S9). This vitamin is essential in many physiological processes (3). Therefore, the possibility to synthetize it *de novo* under low extracellular riboflavin conditions is an advantage. In contrast, clade U has the unique capacity to synthesize *de novo* guanylyl molybdenum cofactor (Supplementary Figure S10), that is essential in molybdenum-dependent enzymes (4). One of such enzymes is the nitrate reductase system Nar. Interestingly, clade U also harbors the operons *nreABC* and *narGHIJ*, that encode for genes that sense anoxic conditions and use nitrate as terminal electron donor instead of oxygen, respectively (5–7). The presence of this molybden-dependent alternative respiratory system exclusively in clade U suggests a different evolutionary history, in which this clade was exposed to anaerobic conditions before adapting to *Drosophila* (Supplementary Figure S11). Similarly, the enzyme sulfur oxidoreductase (EC: 1.8.1.18) is highly enriched in clade C, which suggests that it can potentially use sulfur as terminal electron acceptor in absence of oxygen (Supplementary Figure S11).

Among the genes that were uniquely present in clade C, we found *sbnA* and *sbnB*, that mediate the synthesis of L-2,3-diaminopropionic acid (8). This unusual amino acid serves as precursor of antibiotics and siderophores (8), but the rest of the biosynthesis pathways are absent in the genomes. However, by itself is also a potent enzymatic inhibitor (9,10). Although it goes beyond the scope of this work, we hypothesize that the synthesis of this compound by clade C could be responsible for the observed dose-dependent toxicity upon inoculation in the axenic host (Figure 6). As a sanity check, we blasted the two protein sequences against the clustered non-redundant NCBI database. The best matches had identities of maximum 60%, and belonged to *Streptococcus*, *Bacillus* and *Chitinophaga*. No publicly available *L. plantarum* encodes these genes, which further supports the exceptionality of clade C observed in the phylogeny (Figure 3).

---

**Figure S1.**
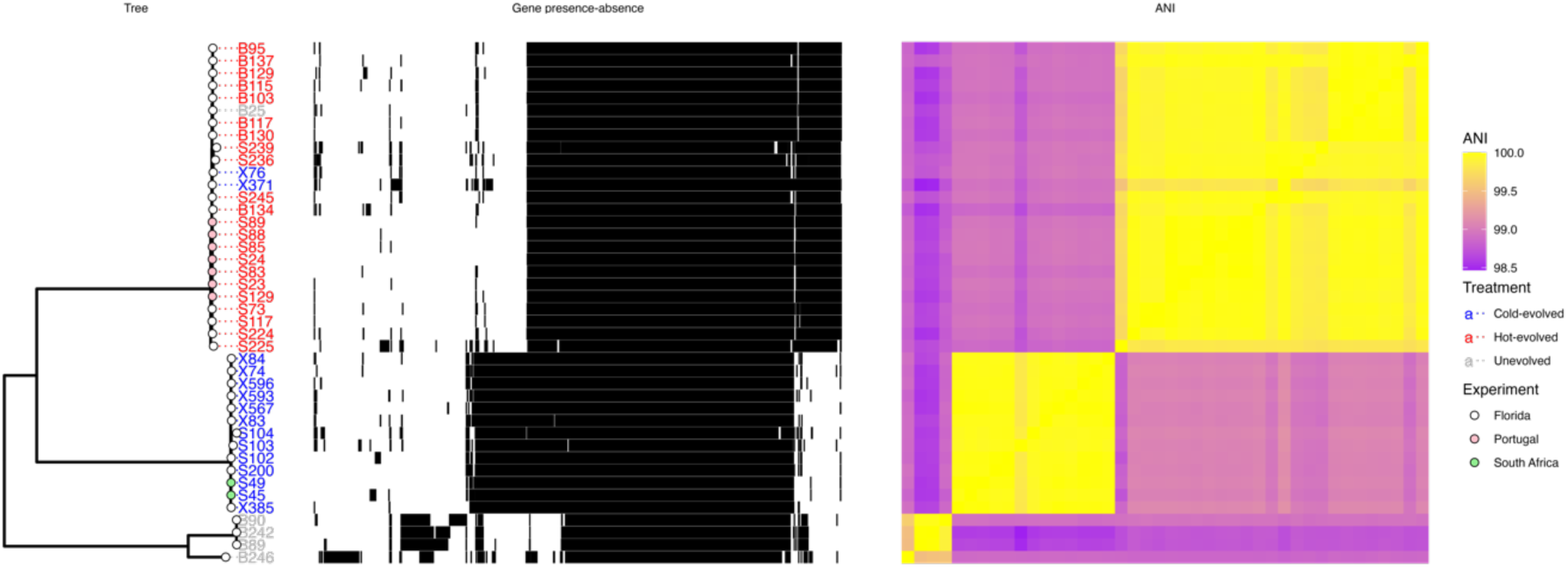
Pangenome of all *L. plantarum* genomes (Table S1), sorted by phylogeny. Left panel: Maximum likelihood tree built based on the core genome alignment. Tree leaves correspond to the name of the isolate and are coloured based on isolation origin: unevolved populations (grey), hot-evolved populations (red) or cold-evolved populations (blue). Tree tips are coloured based on the experiment from which the isolate was obtained: Florida (white), Portugal (pink) or South Africa (green). Middle panel: Pattern of genes presence (black)/absence (white) in each genome. Right panel: heat map depicting the pairwise average nucleotide identity between the isolates. The colour ranges between 98.5% (purple) and 100% (yellow).

**Figure S2.**
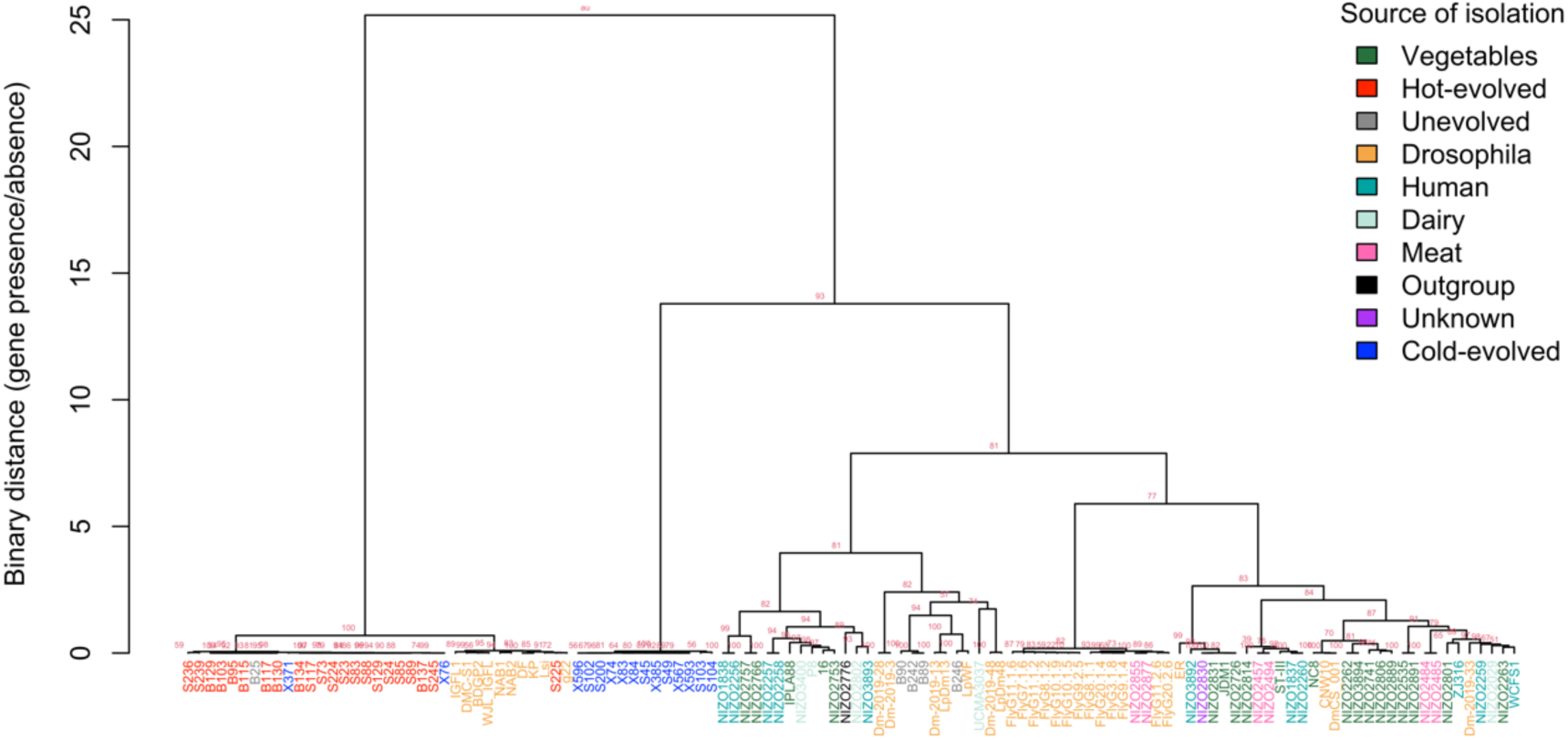
Hierarchical clustering of 92 *L. plantarum* genomes based on the presence-absence of accessory genes. Tree leaves are coloured by source of isolation. The genomes from our lab are coloured according to experimental treatment: unevolved in grey, hot-evolved in red, and cold-evolved in blue. Red values in each node indicate the p-value for each cluster’s robustness based on 1000 bootstrap iterations.

**Figure S3.**
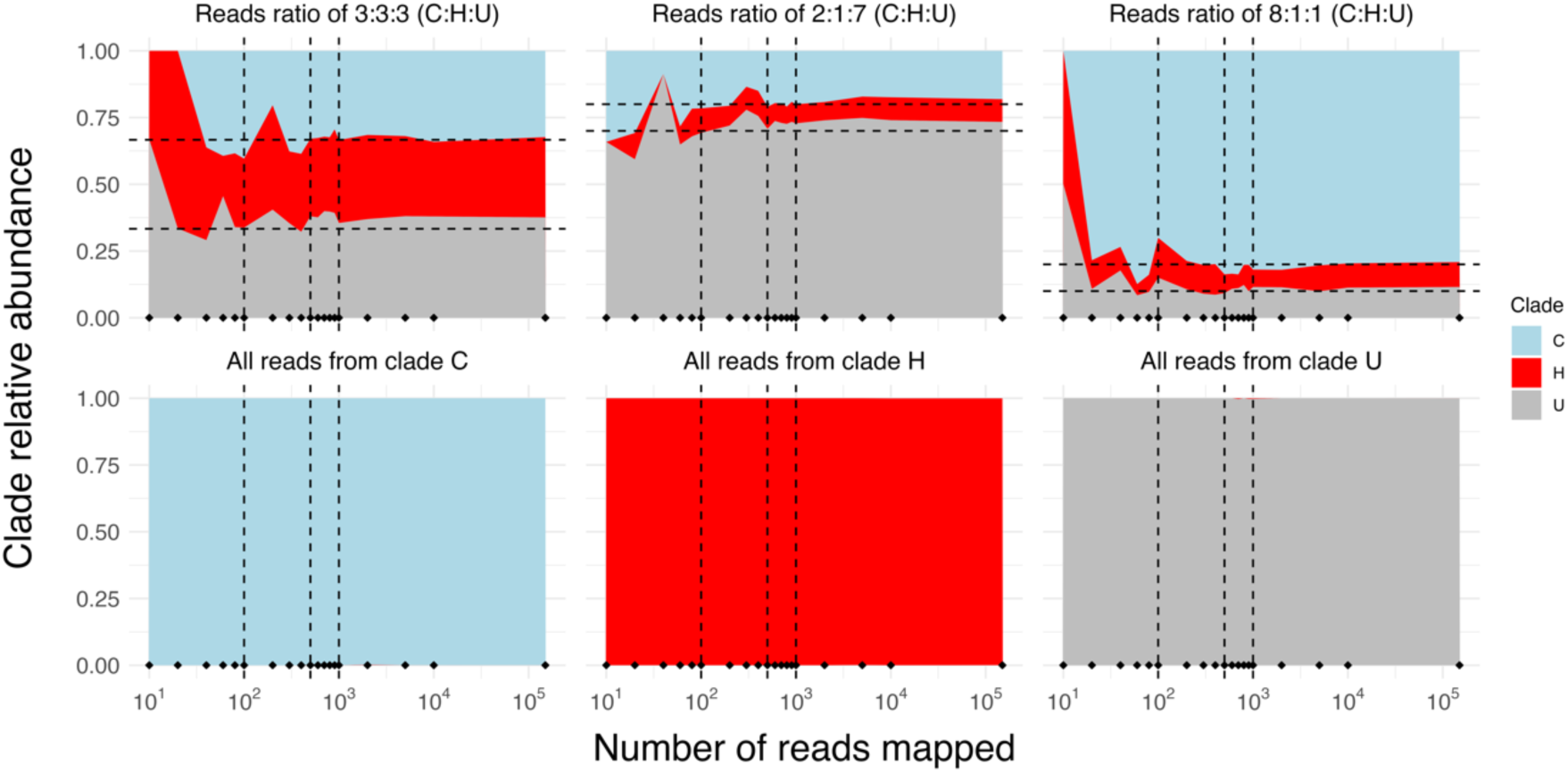
Relative abundance of each *L. plantarum* clade across different reads depths in simulated reads sets with known clade composition. Dashed vertical lines show the values of 100, 500, and 1000 reads. Dashed horizontal lines show the true simulated relative abundances of each clade in the reads set. The top row titles reference to the ratio of reads from each clade following descending order: C:H:U. The bottom plots depict scenarios in which all the reads belong to a single taxon. Diamonds at the bottom of each panel show the sampled number of reads. The x-axis was log10-scaled to show better the sampling distribution.

**Figure S4.**
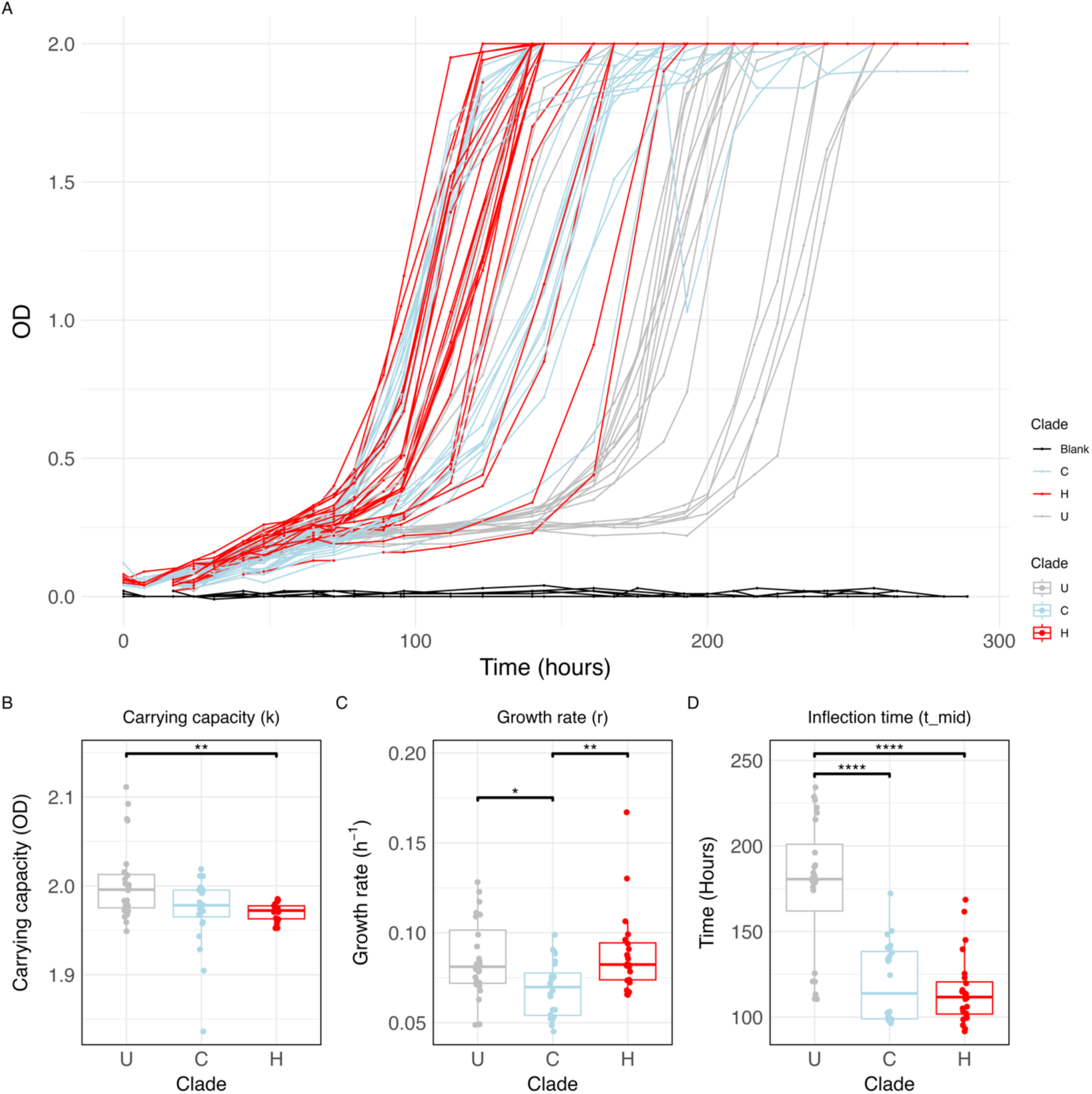
Clade-specific growth dynamics at constant 20 °C. (A) Overlapped curves of all replicates. Lines are coloured by clade. (B-D) Boxplots depicting the carrying capacity, growth rate, and inflection time of each clade. We measured the growth of four isolates from each clade (Supplementary table S1). Each isolate was grown six times, at two different starting points (See Materials and Methods). Each dot corresponds to a technical replicate. Data is grouped and coloured by clade. Statistical significance was determined using Dunn’s test with Holm-adjusted p-values. Only significant comparisons are indicated. **** *p* < 0.0001; *** *p* < 0.001; ** *p* < 0.01; * *p* < 0.05.

**Figure S5.**
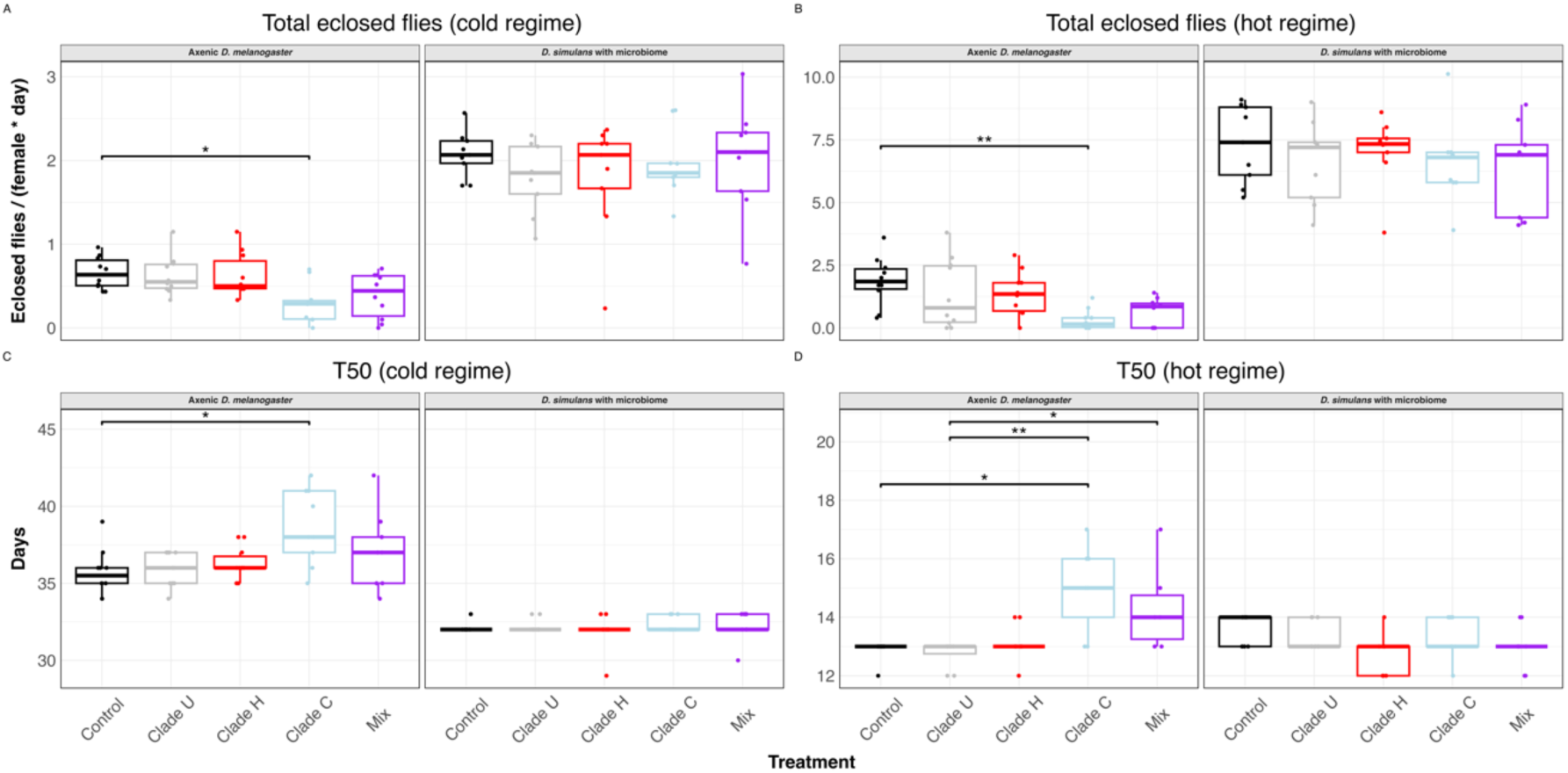
Fitness effect of *L. plantarum* inoculation in axenic *D. melanogaster* and conventionally reared *D. simulans* in the second transfer (See Materials and methods). (A, B) Total number of F1 flies eclosed normalized by day and female under the cold (A) and hot (B) regime. (C, D) Developmental time, estimated as the number of days it takes 50% of the offspring to eclose. Measurements were grouped by inoculation treatment and *Drosophila* species. Each dot corresponds to a biological replicate (n = 10 for *D. melanogaster* and n = 9 for *D. simulans*). Statistical significance was determined using Dunn’s test with Holm-adjusted p-values. Only significant comparisons are indicated. ** p < 0.01; * p < 0.05.

**Figure S6.**
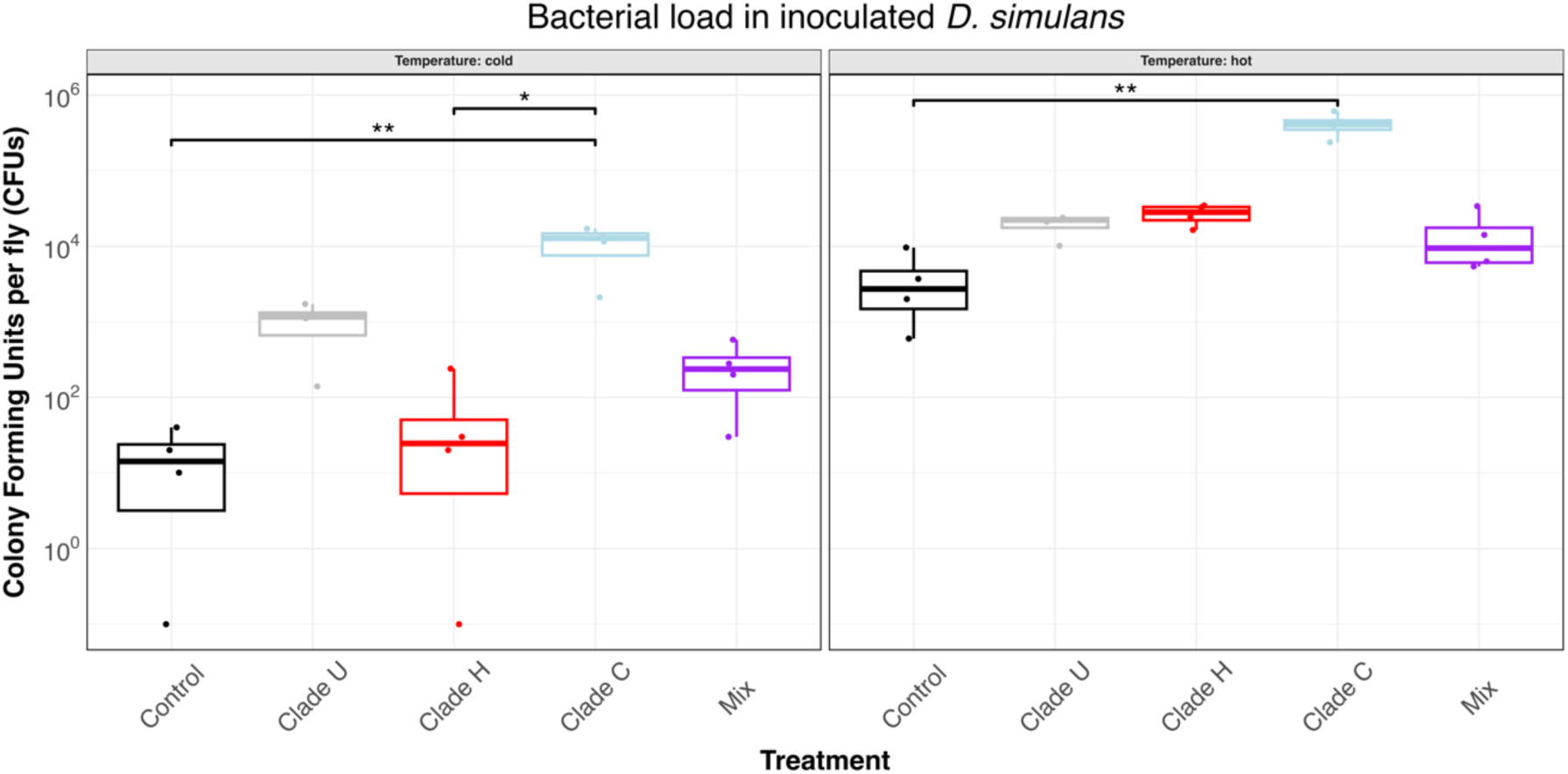
Bacterial load of *D. simulans* individual flies inoculated with *L. plantarum*. Four individual flies were allowed to feed in the corresponding inoculum for 24 h, and then moved to axenic vials for 72 h before crunching and plating their content. Measurements were grouped by inoculation treatment. Each dot corresponds to an individual fly (n = 4). A pseudo-count of 0.1 was added to fit zeros in the log-scaled y-axis. Statistical significance was determined using Dunn’s test with Holm-adjusted p-values. Only significant comparisons are indicated. ** p < 0.01; * p < 0.05.

**Figure S7.**
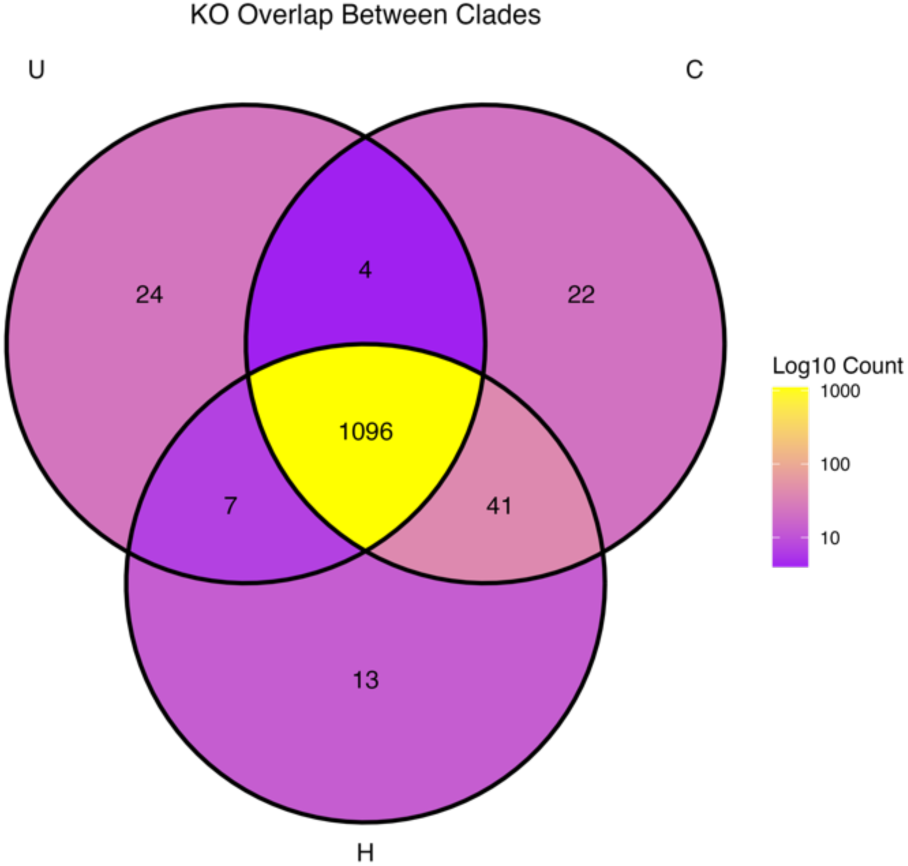
Venn diagram depicting the overlap in KEGG Orthologs between the three clades. Segments were coloured by number of KOs.

**Figure S8.**
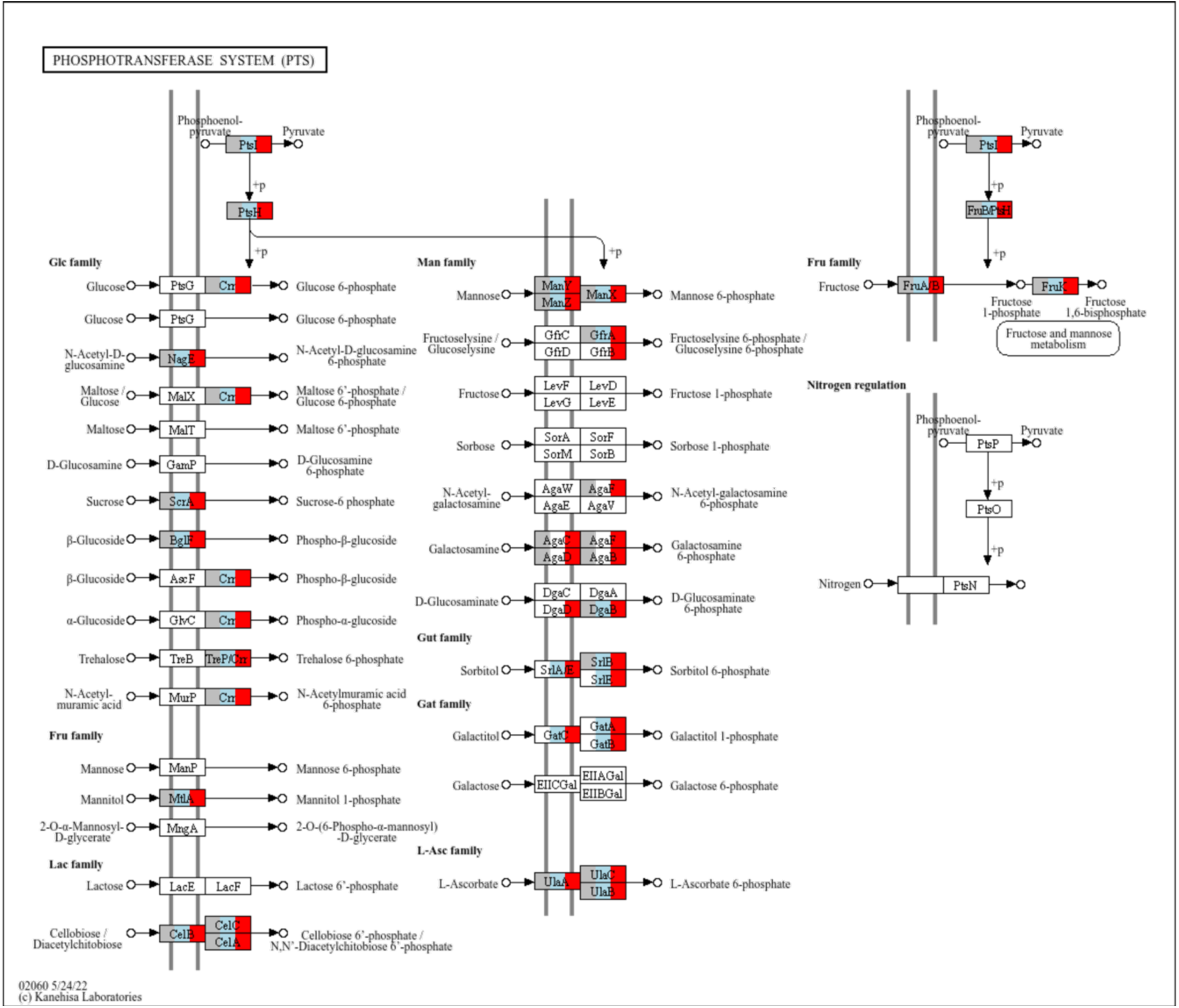
Graphical representation of the KEGG pathway map02060 (Phosphotransferase system). Each rectangle represents a KEGG Ortholog (KO). Orthologs that are enriched in each clade are coloured in grey (present in clade U), blue (present in clade C) and/or red (present in clade H). The three clades differ in the set of sugar-related PTS transporters. Clades C and H have the capacity to internalize sorbitol and galactitol, whereas clades H and U have the ability to import galactosamine.

**Figure S9.**
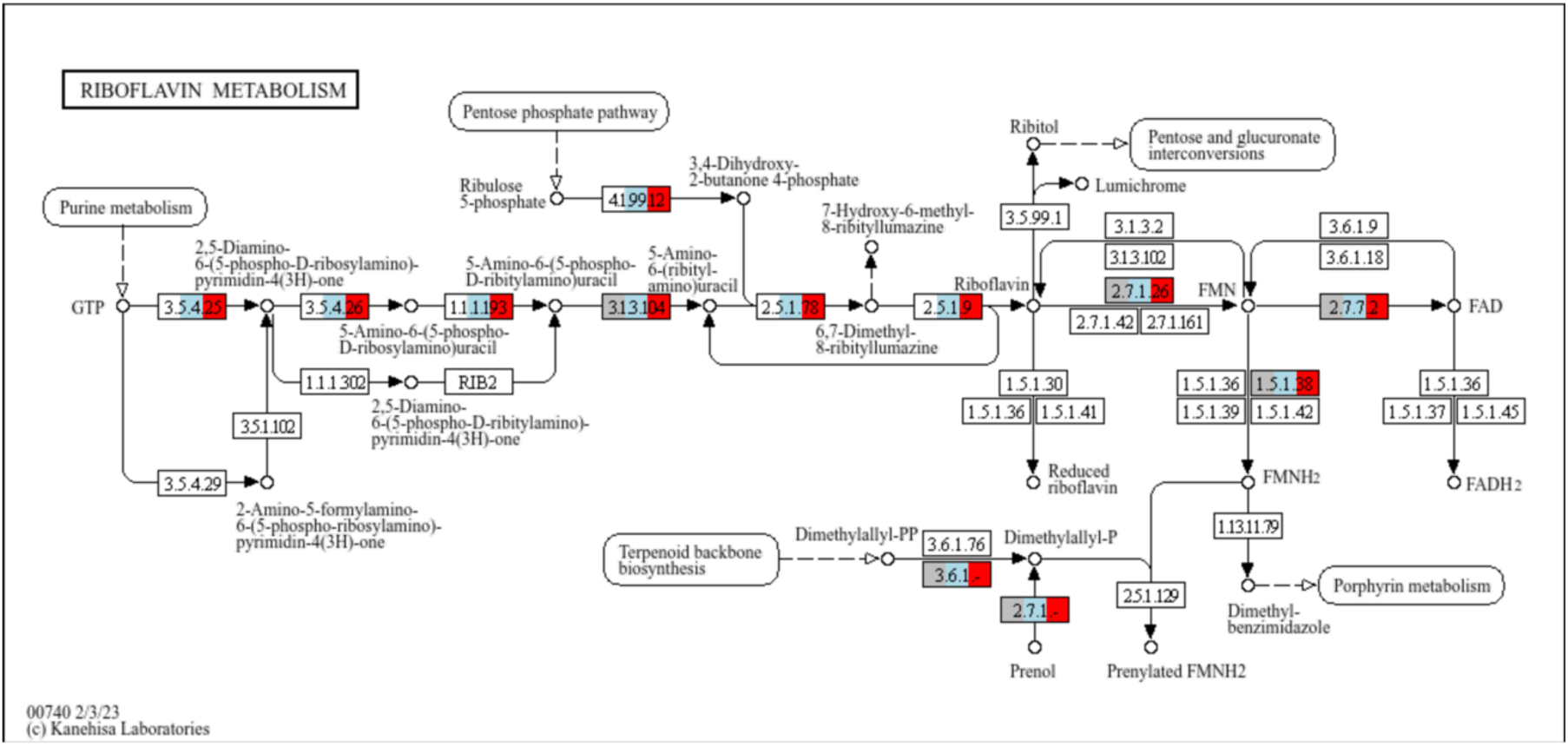
Graphical representation of the KEGG pathway map00740 (Riboflavin metabolism). Each rectangle represents a KEGG Ortholog (KO). Orthologs that are enriched in each clade are coloured in grey (present in clade U), blue (present in clade C) and/or red (present in clade H). Clade U lacks the capacity to produce de novo FAD from GTP, since it is lacking several genes from the pathway.

**Figure S10.**
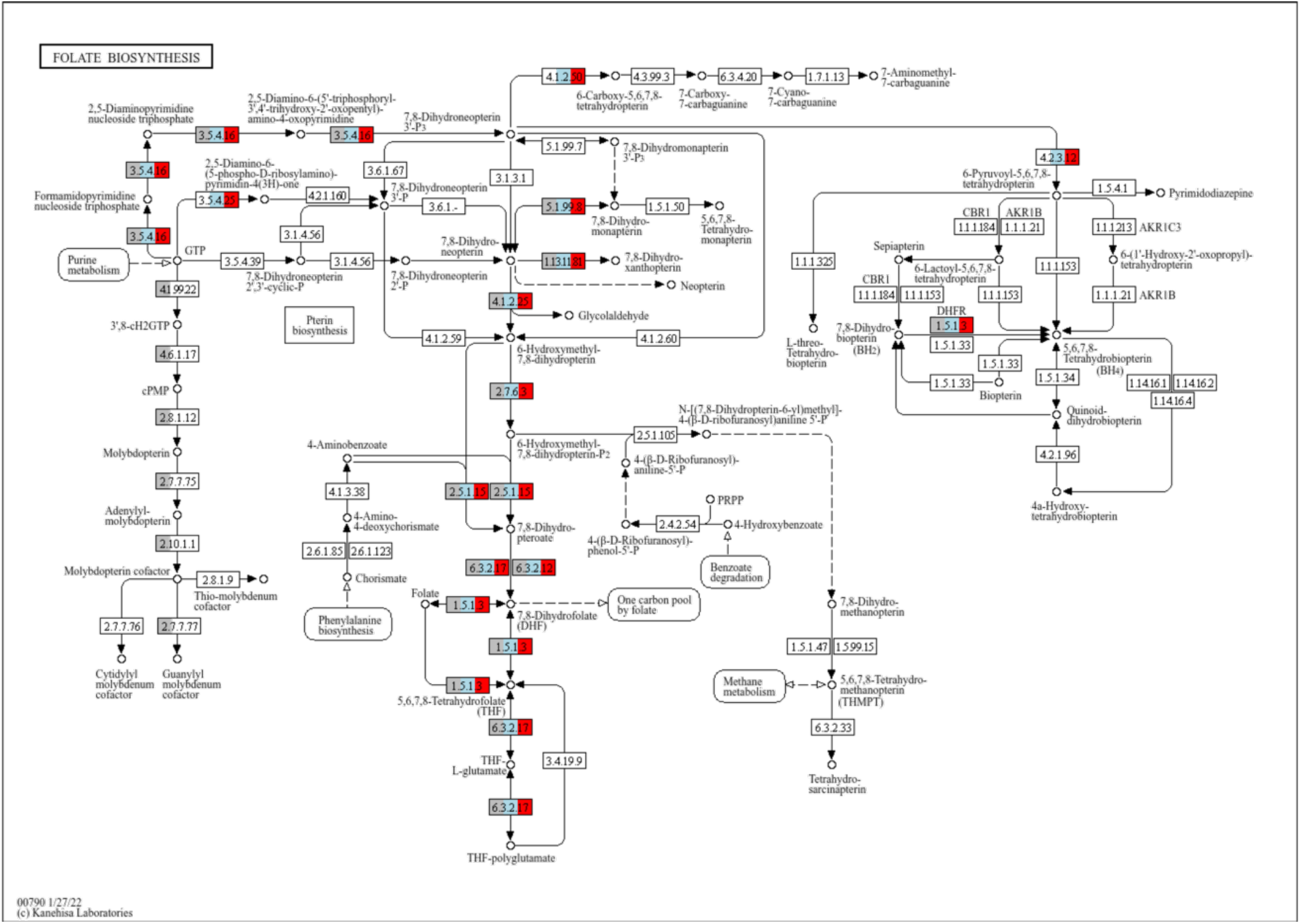
Graphical representation of the KEGG pathway map00790 (Folate biosynthesis). Each rectangle represents a KEGG Ortholog (KO). Orthologs that are enriched in each clade are coloured in grey (present in clade U), blue (present in clade C) and/or red (present in clade H). Clade U has the unique capacity to synthesize de novo guanylyl molybdenum cofactor.

**Figure S11.**
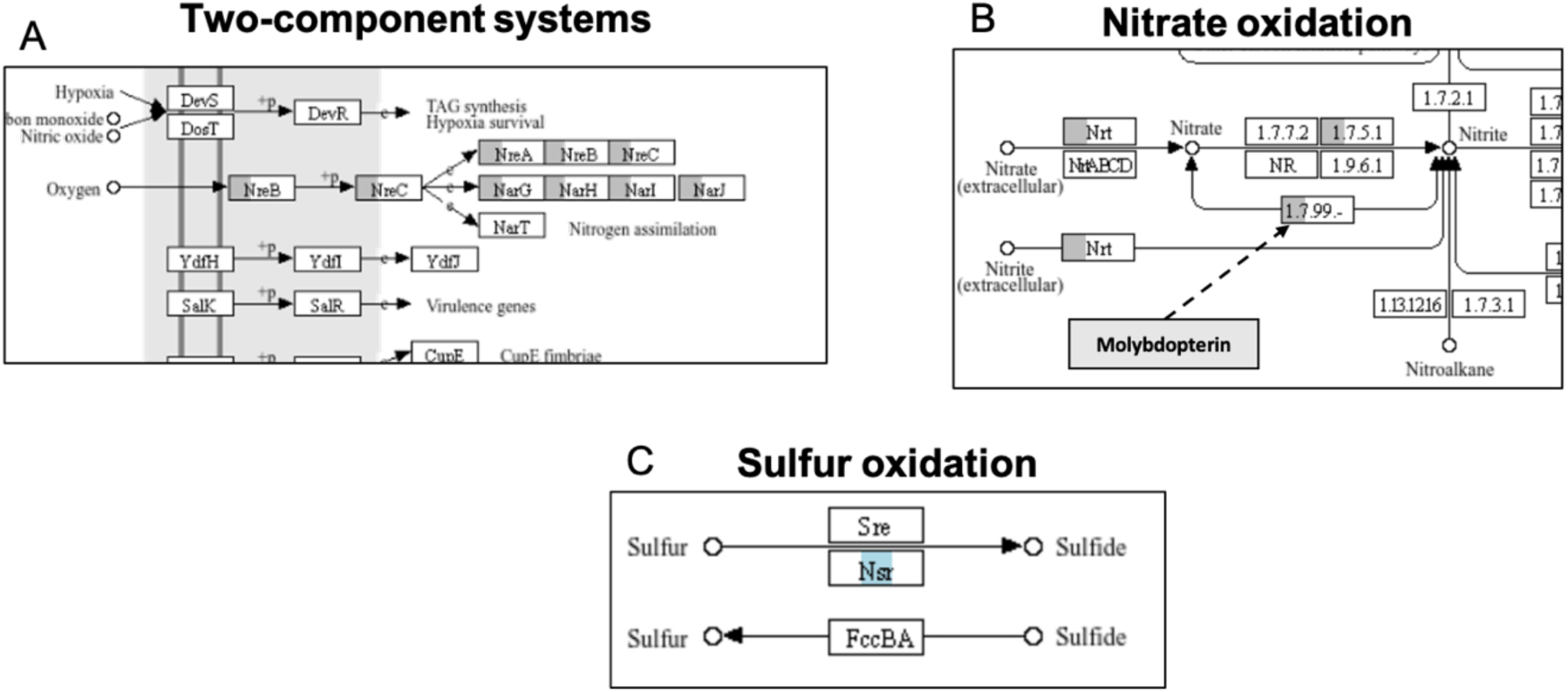
Subset of the KEGG pathways map02020 (Two-component system; A), map00910 (Nitrogen metabolism; B), and map00920 (Sulfur metabolism; C). Each rectangle represents a KEGG Ortholog (KO). Orthologs that are enriched in each clade are coloured in grey (present in clade U), blue (present in clade C) and/or red (present in clade H). Clade U encodes a unique alternative respiratory system. The operons nreABC and narGHIJ encode for genes that sense anoxic conditions and use nitrate as terminal electron donor instead of oxygen (A, B). The enzyme sulfur oxidoreductase (Nsr) is highly enriched in the clade C, which suggests that it can potentially use sulfur as terminal electron acceptor in absence of oxygen (C).

**Figure S12.**
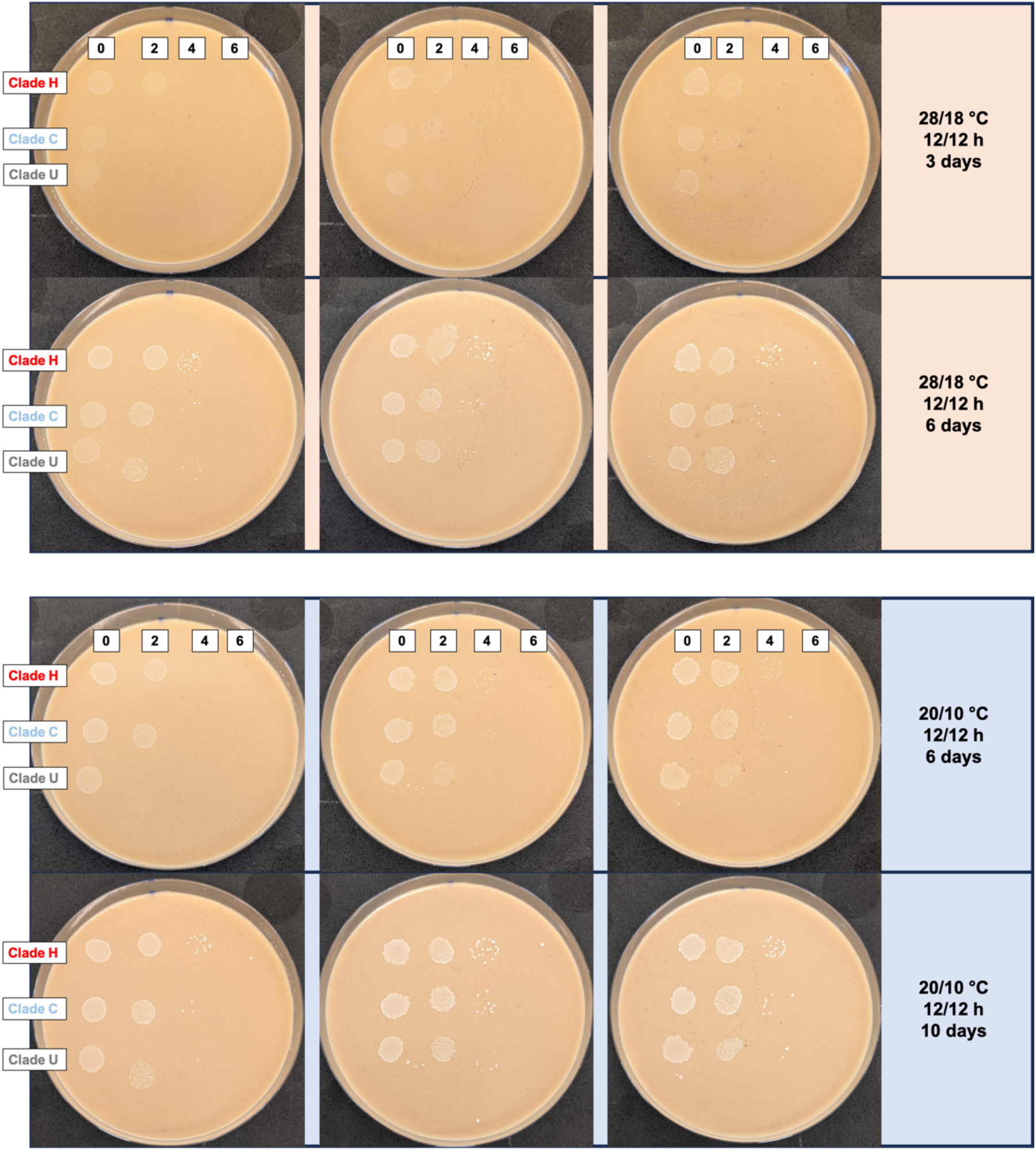
We quantified bacterial growth speed in solid fly food in an agar-drop assay (See Materials and Methods). Briefly, six plates were cultivated with a 3x4 grid of 10 µl drops from each clade (rows) at four concentrations (columns); from the original suspension (0) to dilution 1:10^6^ (6). Three plates were incubated at either hot (top panel) or cold (bottom panel) temperature conditions. We recorded growth progression after incubation days three and six for the hot, and six and ten for the cold. Regardless of the temperature regime, we observed delayed growth of clade U relative to clades C and H. This was seen as overall dimmer drops of this clade in dilution 2 in the first time point (top sub-panels). In the second time point, most drops had reached bacterial saturation (bottom sub-panels).

## Tables

**Table S1 (separate file).** Samples information and data availability of the isolates generated in the study.

**Table S2 (separate file).** Samples information and data availability of the publicly available genomes used to build the *L. plantarum* phylogeny.

**Table S3 (separate file).** Biologically relevant clade-enriched functions based on the KEGG enrichment analysis.

**Table S4 (separate file).** List of taxa isolated from the experimentally-evolved flies.

**Table S5 (separate file).** Data availability for Pool-Seq samples used in the study.

## Notes

### Competing Interest Statement

The authors have declared no competing interest.

### Summary of Updates

Incorporating reviewers' comments, which resulted in some changes in the text and figures

